# Loss of HNF1B drives pancreatic Intraductal Papillary and Mucinous Neoplasms (IPMN) initiation

**DOI:** 10.1101/2025.06.12.659269

**Authors:** Diane Lorenzo, Lina Aguilera Munoz, Louis Marstrand-Daucé, Anaïs Chassac, Pascal Nicole, Lingkang Meng, Laurence Heidet, Bertrand Knebelmann, Camille Pignolet, Sabrina Doblas, Alain Couvineau, Irene Esposito, Vinciane Rebours, Rémy Nicolle, Jérôme Cros, Anne Couvelard, Cécile Haumaitre

## Abstract

**Background:** Intraductal papillary mucinous neoplasms (IPMNs) are clinically detectable precursors of pancreatic adenocarcinoma, yet the mechanisms initiating their development remain poorly defined. Although KRAS mutations are highly frequent in human IPMNs, KRAS activation in pancreatic ductal cells alone fails to recapitulate IPMN development in murine models, indicating that additional tumor-suppressive mechanisms must be overcome.

**Objective:** The objective was to determine whether loss of the transcription factor HNF1B predisposes to initiation of IPMN.

**Design:** We assessed HNF1B nuclear expression and promoter methylation in resected human IPMN specimens. To model IPMN initiation, we generated mice with ductal-specific inactivation of *Hnf1b*, alone or combined with KRAS^G12D^. Ductal organoids and RNA-sequencing were used to investigate molecular mechanisms. Transcriptomic analyses were also performed on human IPMN surgical specimens. MRI from germline *HNF1B* mutation/deletion carriers was re-evaluated for IPMN prevalence.

**Results:** Human IPMNs showed loss of HNF1B by immunochemistry, with enrichment to promoter methylation that increased with dysplasia grade. The KHC model recapitulated the key features of IPMN development including ductal dilation, high proliferation, papillary architecture and mucin production. Loss of Hnf1b together with Kras activation induced loss of primary cilia, cellular reprogramming and engaged oncogenic YAP and Wnt/β-catenin signaling, similar to human IPMNs. Moreover, germline *HNF1B* carriers exhibited a markedly increased prevalence of branch-duct IPMN.

**Conclusion:** HNF1B functions as a tumor-suppressive gatekeeper of pancreatic ductal cells. These findings highlight *HNF1B* inactivation as a potential biomarker and therapeutic entry point for early interception of IPMN-driven pancreatic cancer. They also have implications for the surveillance of HNF1B-syndrome.

## Introduction

Pancreatic ductal adenocarcinoma (PDAC) remains one of the deadliest cancers, with a 5-year survival of less than 10% [1]. This poor prognosis reflects late detection and limited therapeutic options [1]. Understanding the mechanisms of tumorigenesis is critical for early detection and improving outcomes.

PDAC arises primarily from two preneoplastic lesions: Pancreatic Intraepithelial Neoplasia (PanIN) and Intraductal Papillary Mucinous Neoplasm (IPMN) [2–4]. Mouse models with lineage tracing have demonstrated that PanIN mainly originate from acinar cells and IPMNs from ductal cells [5]. IPMNs are defined by papillary growth of pancreatic ductal cells with mucin secretion. They are subclassified histologically as gastric, intestinal or pancreatobiliary according to morphology and mucin immunostaining [6]. Unlike PanINs, which are microscopic and undetectable by imaging, IPMNs are clinically visible as cystic dilations of pancreatic ducts and represent the only screen-detectable precursor lesion [6]. IPMNs affect 5–10% of adults, with prevalence increasing with age [6]. Although not all IPMNs progress to cancer, their unpredictable evolution necessitates careful monitoring [6].

The molecular mechanisms driving the initiation and progression of IPMNs remain incompletely understood. Genomic studies have shown that the most frequently mutated genes in IPMNs are KRAS, GNAS and RNF43 [7–10]. Although KRAS mutations are present in 90% of IPMNs [7,8,11], activating KRAS^G12D^ in pancreatic ducts of genetically engineered mouse models (GEMMs) has failed to induce IPMNs, even under inflammatory conditions [2]. This indicates that additional genetic or epigenetic changes are necessary. While such GEMMs exist, most of them induced mutations in pancreatic progenitors rather than in ducts, implying that they do not fully recapitulate human IPMN development [3,12,13]. Studies on GEMMs have implicated PTEN loss, LKB1 loss and perturbation of epigenetic regulators, as possible drivers of IPMNs [3,13,14]. However, in human IPMNs, these alterations are detected in only 5 to 20% of cases, suggesting the existence of additional initiating mechanisms [9,10]. This highlights the need to interrogate ductal-specific drivers of IPMN initiation. Notably, HNF1B, essential for pancreatic development and postnatally restricted to ductal cells, may play an important role in IPMN initiation [15,16]. We previously showed that postnatal ductal-specific inactivation of *Hnf1b* leads to cystic ducts, ductal polarity defects and increased cell proliferation in mice [17]. Across cancers, HNF1B dysregulation can be tumor-suppressive or oncogenic depending on the organ and tumor subtype [18]. Moreover, *HNF1B* was identified as a susceptibility locus for pancreatic cancer through Genome-Wide Association Studies (GWAS) [19].

Accordingly, we set out to determine whether *HNF1B* inactivation contributes causally to IPMN initiation in ductal epithelium. To this end, we triangulated evidence across human IPMN tissues, ductal-specific mouse models and organoids, and a cohort of *HNF1B* germline carriers.

## Materials and Methods

### Immunohistochemical detection of HNF1B in human sporadic IPMN tissues

FFPE tissue sections from surgical specimens (2012-2022) of human IPMNs, were selected on the basis of histopathological subtyping (Beaujon Hospital). HNF1B IHC was performed using the automated Leica3 IHC slide staining system to assess the nuclear expression of HNF1B (antibodies: Sigma HPA00202089 and HPA002083) across all IPMN subtypes (n=61).

### HNF1B methylation profiling in human IPMNs

*HNF1B* methylation was analyzed in human pancreatic ducts, LG-IPMNs (n=32), and HG-IPMNs (n=20) of both gastric (n=32) and intestinal (n=20) subtypes. Data were obtained in collaboration with the German Cancer Research Center (I. Esposito) [11]. IDAT files were loaded into ChAMP, and data analysis was processed using the R-Bioconductor package Illumina Infinium HumanMethylation27 assay (Illuminaio). *HNF1B* methylation was assessed using beta values derived from specific *HNF1B* probes relevant to gene expression and potential epigenetic silencing mechanisms, with the results visualized in a heatmap. Methylation levels were categorized using the following thresholds of beta values: Methylated (>0.5), Unmethylated (<0.4), and Intermediate (between 0.4 and 0.5). Complementary information is provided in Supplementary Material 1.

### RNA-seq analysis of human IPMNs

RNA extraction was performed on microdissected Formalin-fixed paraffin-embedded (FFPE) gastric IPMN tissues using the Quick-RNA FFPE Miniprep ZYMO Kit and the Roche Millisect® AVENIO Microdissector. RNA sequencing (SMARTer Kit Takara) was conducted on surgical samples (2017-2022) from patients with gastric IPMNs with low-grade dysplasia (LGD; n=140) and high-grade dysplasia (HGD; n=39) collected in Beaujon Hospital (Clichy, France). The reads were aligned with the reference Ensemblv107_GRCh38.75 (Homo sapiens samples) via STAR, and gene expression counts were obtained via FeatureCount. Differential transcriptome analysis of gastric LG- and HG-IPMNs was performed using IDEP, ShinyGO, DECAFE. Detailed analysis is provided in Supplementary Material 2.

## Mouse models

All animal experiments followed European ethical guidelines and were approved by the Ethics committee C2EA-121 PARIS-NORD and the French Ministry of Research (APAFiS N°34673). The Hnf1b^fl/fl^, Sox9-CreER and R26R^YFP^ mouse lines have been described previously [16,17,20]. The LSL-Kras^G12D^ mouse line was from JAX stock #008179 [21]. Inactivation of *Hnf1b* and activation of KRAS in pancreatic ducts were performed via tamoxifen-induced Cre-mediated recombination under the Sox9 promoter. The different genotypes were Hnf1b^fl/fl^;R26R^YFP^ namely “H” / Sox9-creER;LSL-Kras^G12D^;Hnf1b^+/+^;R26R^YFP^ namely “K” / Sox9-creER;Hnf1b^fl/fl^;R26R^YFP^ namely “HC” and Sox9-creER;LSL-Kras^G12D^;Hnf1b^fl/fl^;R26R^YFP^ namely “KHC”. Tamoxifen was delivered postnatally via maternal milk by injecting nursing mothers (6mg/40g) with tamoxifen (Sigma-T5648) in corn oil (20mg/ml) four times during the litter’s first week.

### Histological, immunohistochemical and immunofluorescence analyses of mouse tissues

Pancreata were fixed in 4% PFA for 24h, dehydrated, paraffin-embedded, and sectioned (3µm; four levels). HES and Alcian blue were used for routine staining. IHC was performed on a Leica automated platform with DAB/haematoxylin and nuclei were counterstained with Hoechst. YFP-positive recombined cells were visualized using an anti-GFP antibody and a green fluorochrome-conjugated secondary antibody. Antibodies are listed in Supplementary Table 1. Main duct diameter was averaged from three measurements on serial sections; branch-duct diameter from the 3 largest branch-ducts per pancreas. Cell proliferation was quantified by Ki67 staining (% positive cells/duct), with ducts outlined in Qupath software.

### Magnetic Resonance Imaging (MRI) analysis in mice

To identify pancreatic ducts of mice at the age of 12 weeks, T2-weighted images (2D-acquisition: multi-slice axial acquisition, in-plane resolution 200µm, slice thickness 600 µm, RARE sequence, echo time 25ms, repetition time 4500ms, 4 averages; and 3D acquisition: thin slices of 250µm, RARE sequence, echo time 25ms, repetition time 2500ms, 2 averages) were acquired on a 7T Bruker PharmaScan MRI system, with a 40mm volume coil for transmission/reception (platform FRIM (UMS34), CRI, University of Paris Cité, INSERM).

### Pancreatic ductal organoid culture and RNA-seq analysis of mouse samples

Pancreata from 12-week-old H (n=5), HC (n=4), and KHC (n=4) mice were collected, chopped into small pieces, and digested with collagenase XI (Sigma-C7657). Pancreatic ducts were sorted from the cell suspension and seeded into 50µl BME2 (Cultrex Basement Membrane Extract, Type 2, Bio-Techne, 3532-010-02) matrix with specific culture medium. Organoids were cultured for 7 days. On day 7, organoids were fixed for histology or processed for RNA extraction (RNeasy Micro Kit, Qiagen 74004). The detailed protocol for organoid culture is provided in Supplementary Material 3. RNA library preparation was performed using SMARTer Stranded Total RNA-Seq Kitv3 - Pico Input Mammalian. cDNA and final libraries were controlled with AGILENT Tapestation 2200 on High Sensitivity DNA chip. Sequencing was perfomed on a Novaseq X illumina using a 10B flow cell 2×100 read length, aiming for a total of 30M reads per sample. Reads were aligned with the reference genome Ensemblv107_GRCm38.75 (Mus musculus samples) via STAR, and gene expression counts were obtained via FeatureCount. Extensive analysis of the differential expression transcriptome of ductal organoids (H n=5; HC n=4; KHC n=4) was carried out using Integrated Differential Expression and Pathway (IDEP, ShinyGO, DECAFE©). Detailed RNAseq analysis is provided in Supplementary Material 2. The RNA sequencing datasets generated in this study have been deposited in the Gene Expression Omnibus (GEO) database under accession code GSE298215. https://www.ncbi.nlm.nih.gov/geo/query/acc.cgi?acc=GSE298215.

### Patient cohort with constitutive *HNF1B* mutations

To identify the presence of IPMN in patients with *HNF1B* germline mutations or deletions, all the cross-sectional images (contrast-enhanced CT-scans, MRI) or endoscopic ultrasounds from patients of the Nephrology departments (Necker Hospital, Paris, France) were reviewed. Patient imaging studies were re-evaluated to identify those with IPMN, defined as branch-duct IPMN if a communicating cystic lesion greater than 5mm was present, or as main duct IPMN if there was pancreatic duct dilation greater than 5mm without obstruction [6]. This study was approved by the research Ethics Committee Institutional Review Board (IRB00006477).

### Statistical analyses

Statistical analyses were conducted using GraphPad Prism. Data are shown as means ± standard deviation (SD). Statistical significance (p<0.05) was assessed using descriptive statistics, chi-square tests or Fisher’s exact tests for qualitative variables, Student’s t-tests with Welch correction or ANOVA for multiple comparisons. Fisher’s exact test assessed *HNF1B* methylation across IPMN subtypes and grades. IPMN incidence in *HNF1B*-mutant patients was compared to an age-matched population using Poisson regression.

Patients and the public were not involved in the design, conduct or reporting of this study; clinical management followed routine care when indicated.

## RESULTS

### HNF1B is epigenetically downregulated in human IPMNs

While HNF1B is normally localized to the nuclei of pancreatic ductal and centroacinar cells, IHC showed loss of HNF1B staining in IPMN lesions as ductal cells acquired a dysplastic phenotype, from the LGD stage onward and across histological subtypes (gastric LGD n=36; gastric HGD n=5; pancreatobiliary HGD n=6; intestinal LGD n=21; intestinal HGD n=6) (Figure 1A–B). These results suggest a potential role for the loss of HNF1B expression in the initiation of IPMNs.

**Figure 1:**
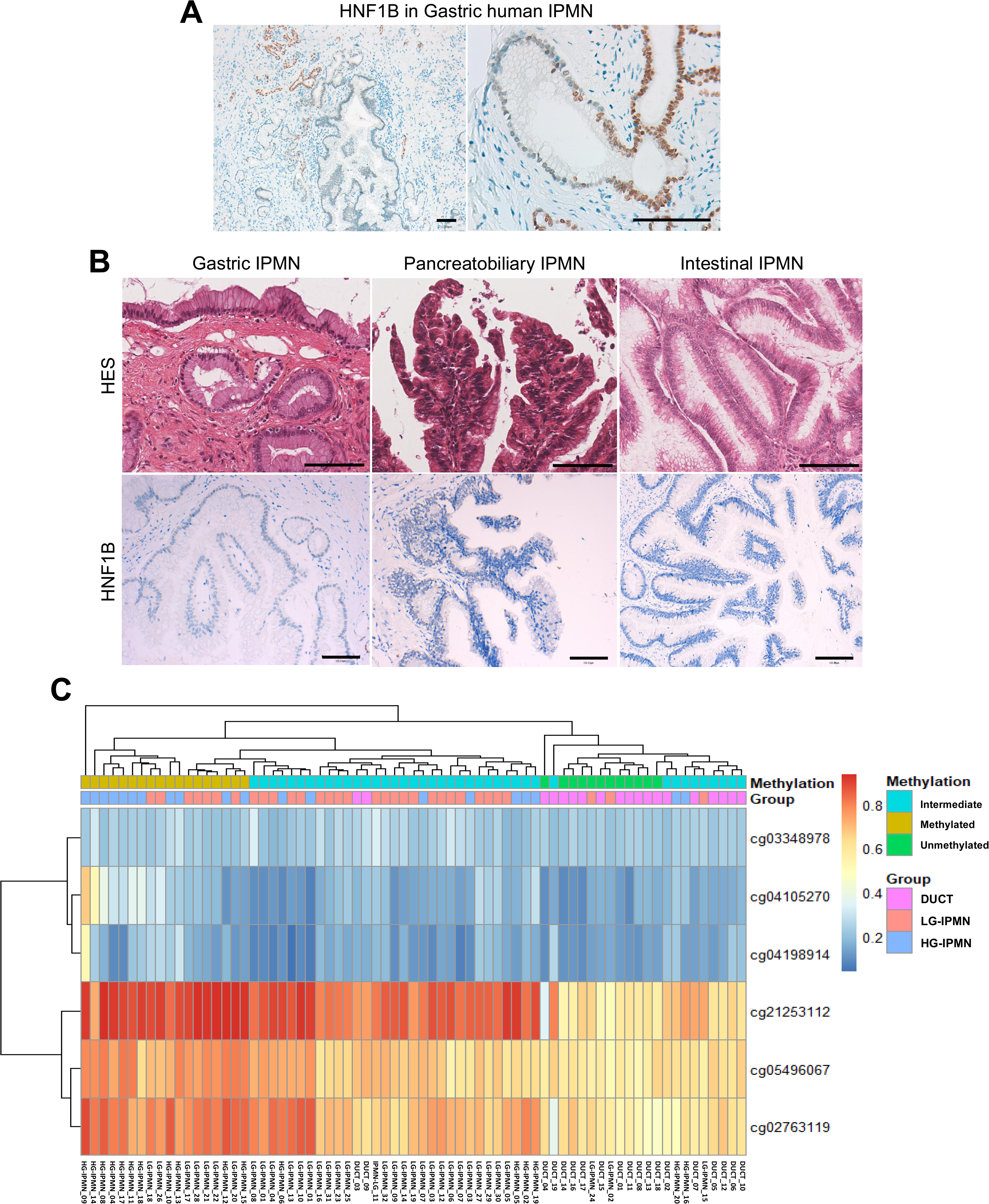
Loss of HNF1B expression in human IPMN. (A) HNF1B immunostaining of human gastric-IPMN showing positive HNF1B staining in ductal cells and negative expression in human gastric IPMN. Right panel: Progressive loss of HNF1B expression with the onset of dysplasia. (B) HES staining and HNF1B immunostaining of human IPMN showing HNF1B extinction in dysplastic cells. (C) Mean CpG methylation of HNF1B probes known to modulate HNF1B expression in human ducts (n=19), gastric (n=32) and intestinal (n=20) IPMN, LG-IPMN (n=32) and HG-IPMN (n=20) (unmethylated: mean β-value <0.4; intermediate: mean β-value >0.4 and <0.5; methylated: mean β-value >0.5). Scale bar, 100 µm.

As *HNF1B* mutations have not been commonly reported in human sporadic IPMN, we explored promoter methylation as a potential mechanism of gene silencing. The *HNF1B* gene had a significantly higher methylation frequency among the 52 IPMNs (unmethylated: 2 (3%), intermediate: 32 (62%) and methylated: 18 (35%)) than among the 19 normal pancreatic ducts (unmethylated: 10 (53%) and intermediate: 9 (47%) (p=1.1×10^-6^)) (Figure 1C). Methylation also increased with grade (LGD n=32: 72% intermediate and 22% methylated; HGD n=20: 45% intermediate and 55% methylated; p=0.037) and did not differ by gastric vs intestinal subtype (Supplementary Figure 1). These data support epigenetic silencing of HNF1B as an important mechanism in human IPMN, contributing to the initiation and progression of dysplastic changes in pancreatic ductal cells.

### Ductal HNF1B loss cooperates with KRAS activation to induce IPMN in mice

To test if *Hnf1b* loss drives IPMN initiation, a GEMM combining *Hnf1b* inactivation and KRAS activation in pancreatic ducts (KHC line: Sox9-creER;LSL-Kras^G12D^;Hnf1b^fl/fl^; R26R^YFP^) was generated. As reported, the pancreatic ducts of K mice (Sox9-creER;LSL-Kras^G12D^;Hnf1b^+/+^;R26R^YFP^) were similar to H controls [3,13]. The pancreatic ducts of HC mice (Sox9-creER;Hnf1b^fl/fl^;R26R^YFP^) were slightly dilated. Phenotypic characterization by MRI, histopathological and IHC analyses demonstrated that KHC mice developed significantly more dilation of the main pancreatic duct (p<0.0001) (Figure 2A-E) and branch ducts (p<0.0001) (Figure 2F-H) compared to controls. Ki67 quantification showed increased ductal cell proliferation in KHC mice compared with controls (HC, K, H) (p<0.0001) (Figure 2I, J).

**Figure 2:**
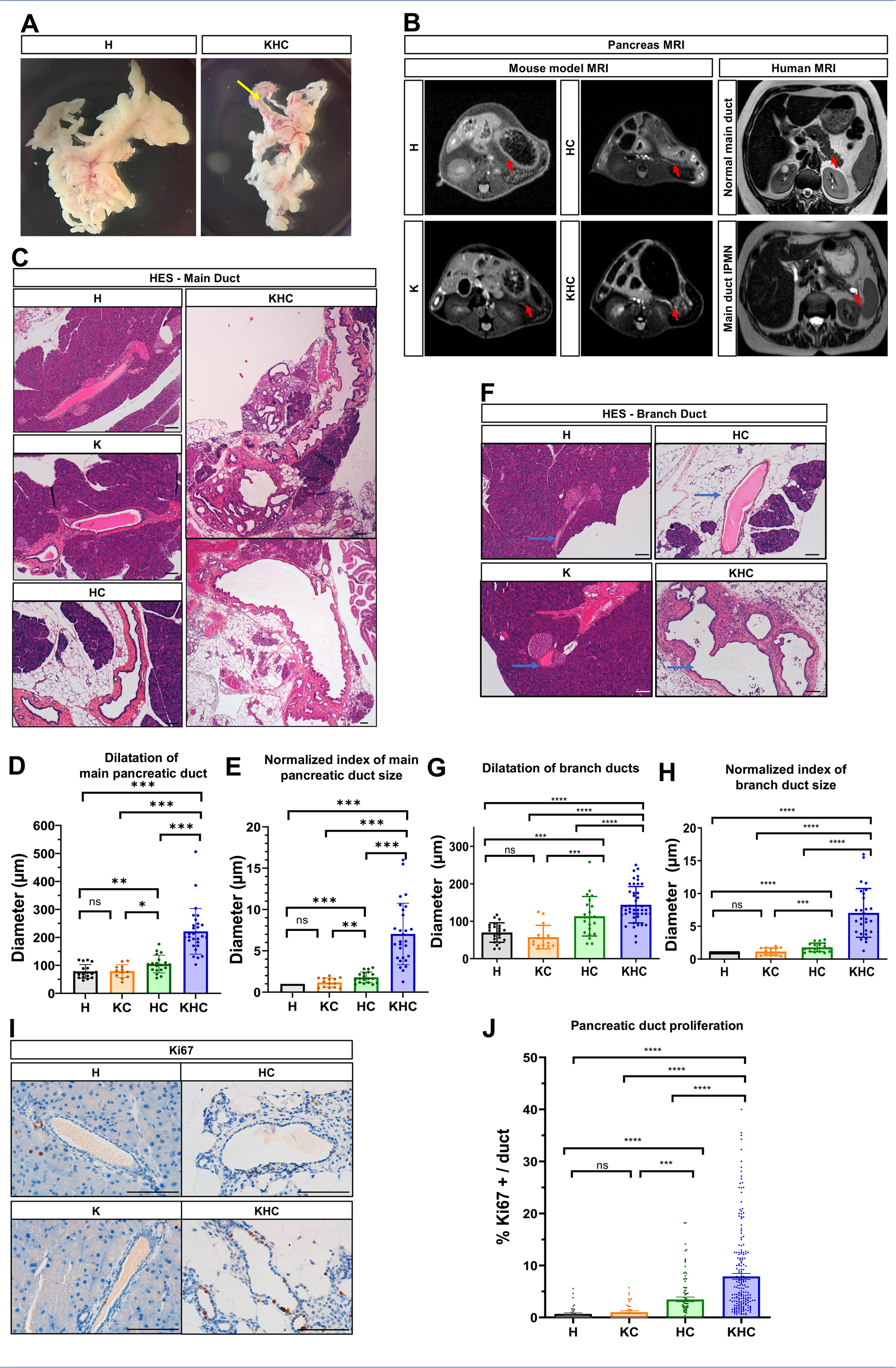
Pancreatic duct morphology and proliferation in KHC vs. controls. (A) Pancreatic tissue from H and KHC mice. KHC mice show main duct dilation (arrow), with reduced parenchyma. (B) T2-weighted MRI of mice (left panels) and human (right panel). T2-weighted MRI: main pancreatic duct is absent in H and K mice, faintly visible in HC, and dilated in KHC. Human images show a normal duct (top right) and a moniliform dilation in the tail consistent with IPMN (bottom right). (C) HES-stained histological section of main pancreatic duct. (D) Diameters of the main pancreatic duct show greater dilation in KHC (n=28) compared to HC (n=19), K (n=12) and H (n=18) mice. (E) Index of main pancreatic duct size (duct dilation normalized to pancreas weight, with control set at 1). The KHC main pancreatic duct was 7 times more dilated than H, and the HC was 2 times more dilated than H. (F) HES-stained histological section of pancreatic branch-duct (blue arrow). (G) Diameters of pancreatic branch-ducts show greater dilation in KHC (n=45) mice compared to HC (n=22), K (n=15) and H (n=22). (H) Index of branch-ducts (branch-duct dilation normalized to pancreas weight, with control set at 1), reveals 7-fold greater dilation in KHC mice compared to H mice, with a 2-fold increase observed in HC compared to H. (I) Ki67 IHC of pancreatic ducts from KHC, HC, K and H mice. (J) Quantification of pancreatic ductal proliferation. Number of ducts counted H (n=43), HC (n=87), K (n=41) and KHC (n=211). * p<0.01, **p<0.001, ***p<0.0001 and ****p<0,00001. Genotypes of the mice: “H” Hnf1b^fl/fl^;R26R^YFP^, “K” Sox9-cre^ER^;LSL-Kras^G12D^;Hnf1b^+/+^;R26R^YFP^, “HC” Sox9-cre^ER^;Hnf1b^fl/fl^;R26R^YFP^, “KHC” Sox9-cre^ER^;LSL-Kras^G12D^;Hnf1b^fl/fl^;R26R^YFP^.

Only KHC ducts displayed dysplasia (p<0.0001) with lesions positive for Alcian blue coloration and Claudin18 IHC, involving both main and branch ducts (Figure 3A-D). KHC lesions displayed papillary/glandular architecture with columnar mucinous (foveolar-like) cells, a morphology typical of gastric-type IPMN (Supplementary Figure 2A), and a gastric immunophenotype (MUC1 apical+/MUC2−) (Supplementary Figure 2B). Critically, IPMN lesions were YFP-positive, including glandular neoplasms, confirming ductal origin (Figure 3E–G’).

**Figure 3:**
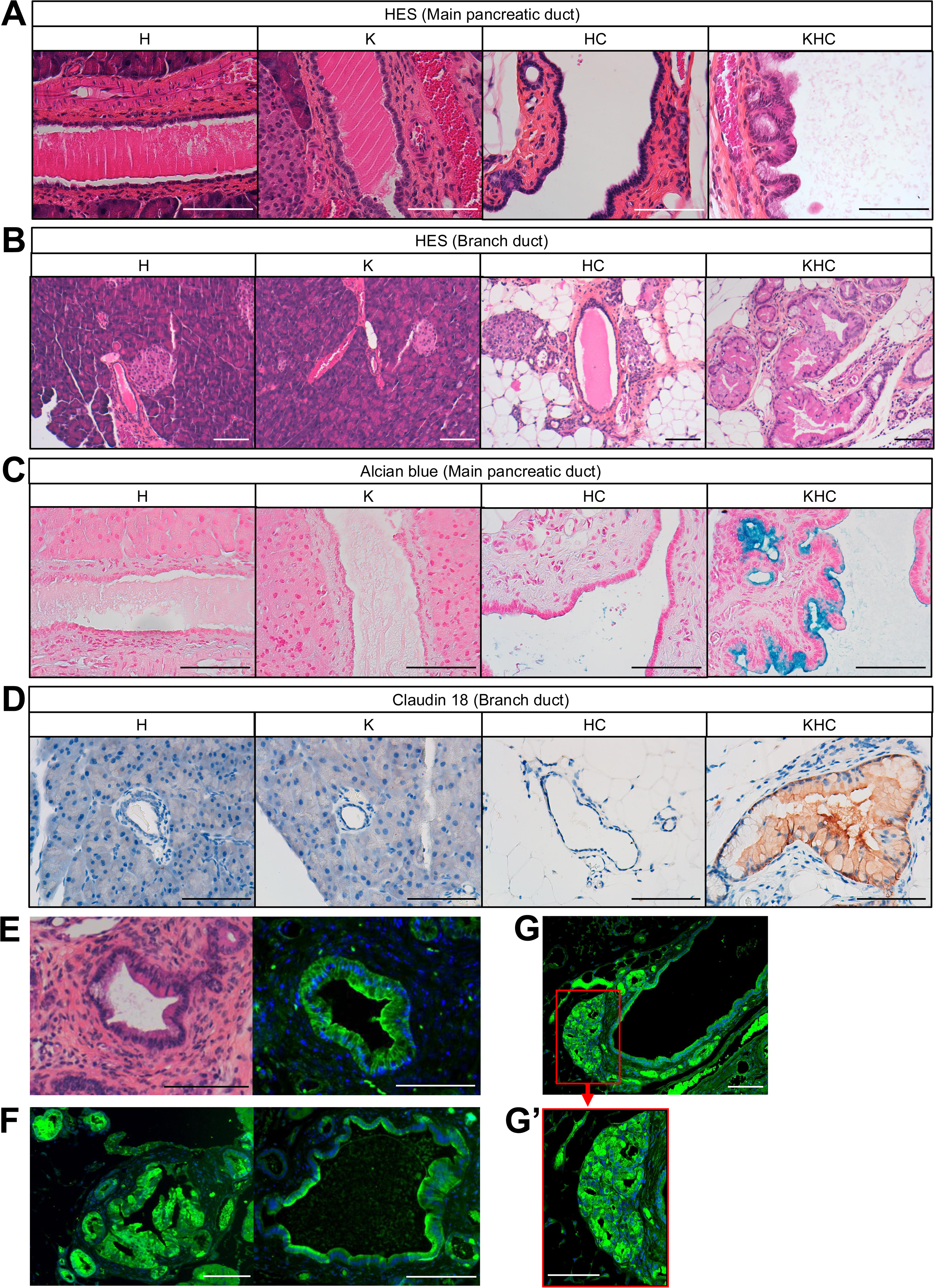
Histological and immunofluorescence characterization of pancreatic ductal lesions in KHC mutants compared to HC, K and H mice. (A) HES staining of the main pancreatic duct of H, K, HC and KHC. (B) HES staining of the branch ducts of H, K, HC and KHC. (C) Alcian blue coloration (mucus presence) of KHC lesion in the main pancreatic duct. (D) Immunohistochemical staining of Claudin-18 showing positive labeling of the KHC lesion in a branch duct. KHC mice exhibit dysplastic lesions with papillary formation, swollen cytoplasm, nuclear deformation, and mucus production, all positively labeled with Claudin 18. (E) HES staining of a ductal dysplasia lesion of a branch duct (left panel) and YFP immunofluorescence (right panel), highlighting IPMN branch-duct cell lineage in KHC mice. Nuclei labeled with Hoechst. (F) Dysplastic papillary lesion of the main pancreatic duct showing YFP-immunofluorescence within dysplasia. (G, G’) YFP immunofluorescence of a main pancreatic duct from KHC mice with an enlarged glandular neoplasm, showing its ductal identity. Nuclei labeled with Hoechst. Scale bar, 100 µm.

Together, these results highlight the synergistic requirement of *Hnf1b* inactivation and KRAS activation to induce IPMN, consistent with a tumor suppressive role of *Hnf1b* in the ductal epithelium.

### Molecular programs associated with *Hnf1b* loss in IPMN initiation

To understand how *Hnf1b* deficiency can lead to IPMN initiation, ductal organoids were cultured 7 days for RNAseq analysis (Figure 4A-B). Growth rates were comparable (H: n=5, HC: n=4, KHC: n=4) (Supplementary Figure 3A), but HC and KHC organoids uniquely formed tubular structures/protrusions, reflecting cell polarity defects resulting from *Hnf1b* deficiency [17] (Figure 4A-B; Supplementary Figure 3B-C). This kind of tubular structure has been previously documented in an *in vitro* model of ducts deficient for *Brg1* [22]. SOX9 immunostaining confirmed the exclusive ductal identity of organoids (Supplementary Figure 3D).

**Figure 4:**
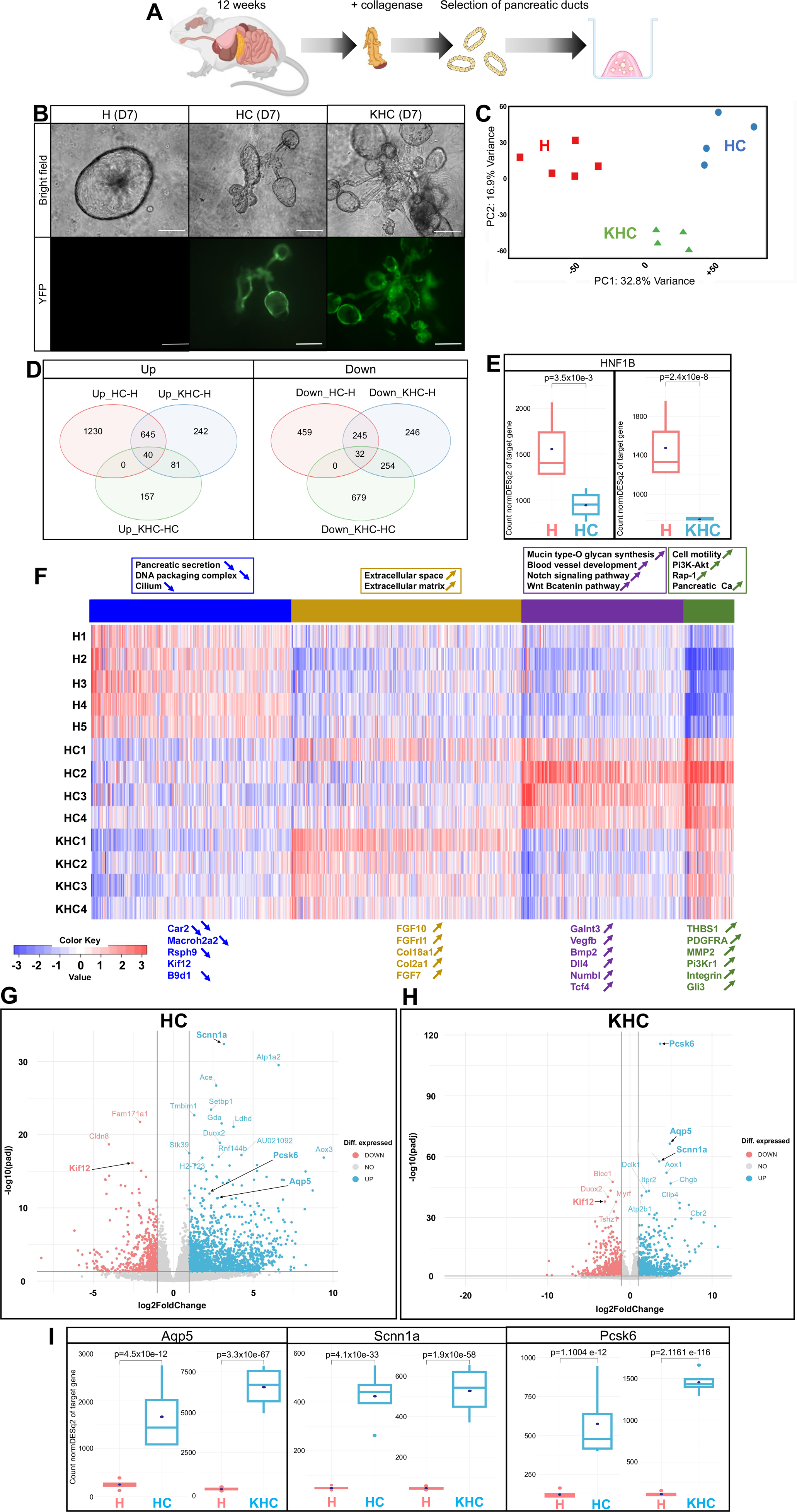
Transcriptomic effects in HC and KHC pancreatic ductal organoids and associated ductal cell dysfunction. (A) Schematic representation of the generation of pancreatic ductal organoids from 12-week-old H, HC, and KHC mice, including pancreatic digestion with collagenase, manual sorting of ductal cells and culture in a BME2 dome for 7 days (created with Biorender). (B) Bright field and green fluorescence (YFP) images of organoids from H, HC, and KHC mice on day 7. Scale bar = 50 µm. (C) Principal component analysis (PCA) of gene expression data across H (red squares, n=5), HC (blue dots, n=4), and KHC (green triangles, n=4) samples. (D) Venn diagrams showing upregulated (left panel) versus downregulated (right panel) genes and their overlap across the following comparisons: HC vs H, KHC vs H, and KHC vs HC. (E) Boxplot showing the transcript levels of Hnf1b in HC and KHC ducts compared with H ducts. (F) K-means heatmap showing clustering of H, HC, and KHC groups, highlighting clusters of downregulated pathways (blue), clusters of upregulated pathways (yellow, purple, green), and examples of associated genes, in HC and KHC compared with control H. (G) Volcano plot, with upregulated genes in blue and downregulated genes in red. Differential gene expression in HC compared with H. (H) Differential gene expression in KHC compared with H. (I) Boxplot showing the upregulation of Aqp5, Scnn1a, and Pcsk6 in HC and KHC compared with H.

Transcriptomic analysis revealed clear distinct H, HC and KHC clusters on unsupervised Principal Component Analysis (PCA) (Figure 4C). The Venn diagram showed that 685 genes were upregulated in HC and KHC compared with the H group, representing 36% of upregulated genes in HC vs H and 68% in KHC vs H. Similarly, 277 genes were downregulated in both, accounting for 38% in HC vs H and 36% in KHC vs H. Overall, many gene changes seen in KHC were already present in HC, with more genes upregulated in KHC than in HC (Figure 4D). As expected, *Hnf1b* transcripts were markedly reduced in HC/KHC ductal cells (Figure 4E).

We next asked which coordinated programs accompany *Hnf1b* loss and the shift toward dysplasia. K-means clustering revealed deregulated gene clusters (Figure 4F). One cluster of downregulated genes in HC and KHC ductal cells, compared to controls, was notably linked to primary cilium function and impaired pancreatic secretion. This included reduced expression of bicarbonate secretion genes, such as *Car2* (p<0.00001; KHC vs H). The heatmap also highlighted upregulation of extracellular matrix genes involved in cell adhesion, migration, and growth factor signaling pathways. To identify key differentially expressed genes in HC and KHC ductal cells, a volcano plot highlighted genes with the most significant expression changes (Figure 4G-H). *Scnn1a* (subunit of sodium channel, ENaC) and *Aqp5* (water channel), a spasmolytic polypeptide-expressing metaplasia (SPEM) marker [23], were among the most significantly upregulated genes (Figure 4I). Both are key regulators of intracellular osmolarity, and their upregulation correlated with reduced bicarbonate secretion; each has also been implicated in oncogenic behavior in pancreatic cancer [23,24]. Moreover, *Pcks6*, linked to tumor proliferation via the Raf-MEK1/2-ERK1/2 pathway [25], was highly upregulated in HC and KHC ductal cells (Figure 4I).

In contrast, *Kif12* was strongly downregulated, reflecting impaired primary-cilium function (Figure 4G, H). Moreover, GOCC and GOBP pathway analyses showed downregulation of cilia-related genes (Figure 5A-B), including *Bbs1*, *Oscp1*, *Kif12*, *Ofd1* and the HNF1B direct targets *Pkhd1* and *Cys1* (Figure 5C; Supplementary Table 2) [26]. Pancreatic ductal cells harbor an immotile primary cilium, a microtubule-based organelle that protrudes from the surface of the cells and acts as a chemo- and mechanosensor; it integrates signaling pathways such as Wnt and Hedgehog [27]. Corroborating these results, immunostaining revealed the dramatic loss of primary cilia in HC and KHC ductal cells (Figure 5D-E).

**Figure 5:**
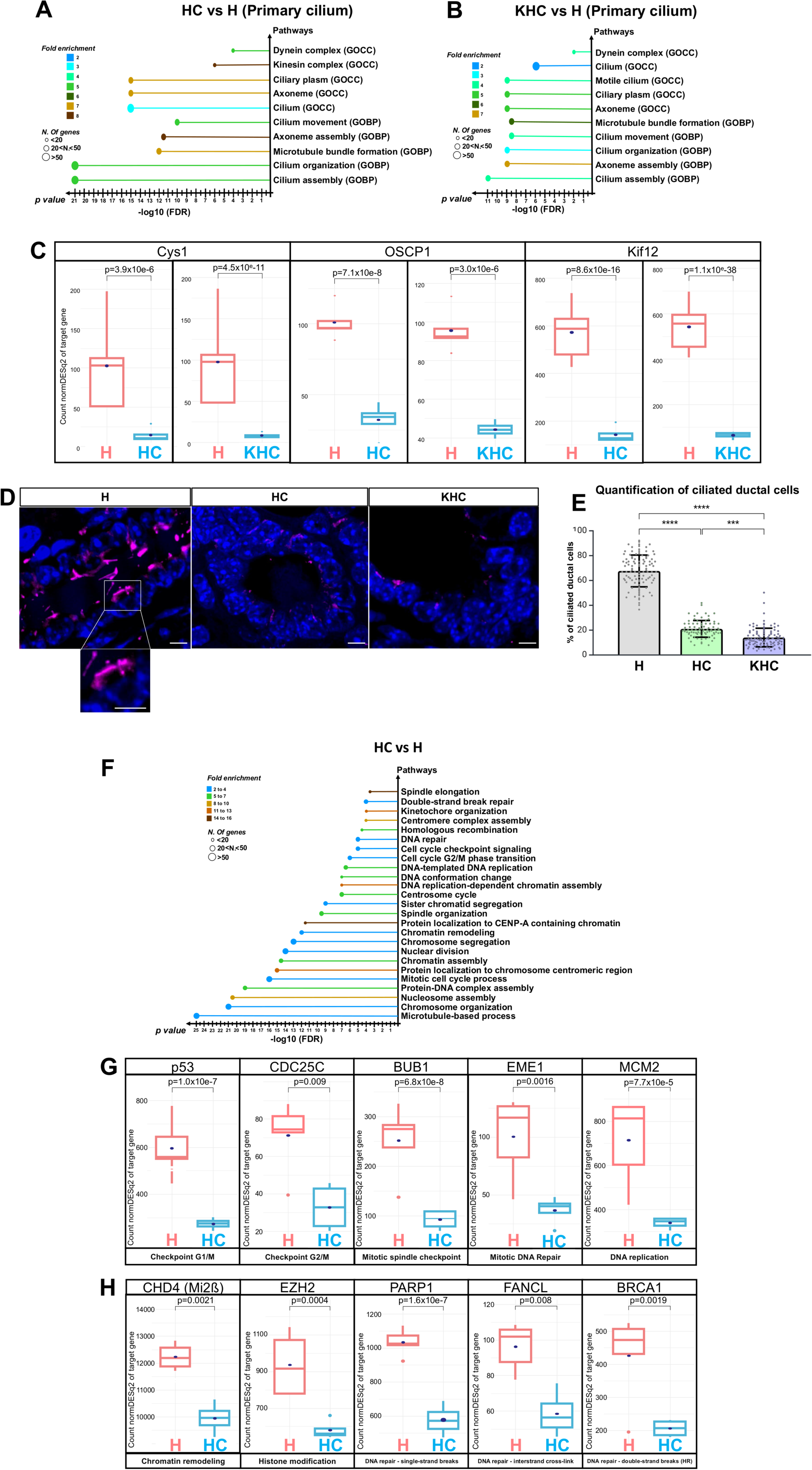
Downregulation of primary cilia-related pathways in HC and KHC, and downregulation of cell cycle/genomic stability regulatory Pathways in HC compared with. **H** (A-B) Lollipop diagrams of the Gene Ontology Biological Process (GOBP) and Gene Ontology Cellular Component (GOCC) pathways related to primary cilia, showing downregulation in HC and KHC ductal cells compared with control H. Each lollipop represents a pathway, with the x-axis indicating the -log10 of the p-value, reflecting the statistical significance of the downregulation, and the y-axis listing the pathway names. The color of the lollipops corresponds to the fold enrichment, with blue indicating lower enrichment and brown indicating higher enrichment. The size of the lollipop dots represents the number of genes involved in each pathway. (C) Boxplots illustrating the downregulation of genes associated with primary cilia (Cys1, Oscp1, Kif12) in HC and KHC ductal cells compared with control H. (D) Immunostaining of acetylated-tubulin (red) showing primary cilia loss in HC and KHC ductal cells. Nuclei are stained with Hoechst. Scale bar: 5 mm. (E) Quantification of ciliated ductal cells (F) Lollipop diagram of downregulated GOBP pathways involved in cell cycle regulation and genomic stability (DNA repair pathways, mitotic spindle, regulation of chromatin structure and function, cell division and its checkpoints) in HC compared with control H. (G, H). Boxplots showing the downregulation of key genes involved in pathways related to (G) cell cycle regulation and (H) genomic stability. HR: homologous recombination

Beyond ciliary and secretory programs, GOBP enrichment also pointed to a coordinated downregulation of pathways governing chromatin structure, cell division, mitotic spindle function, and DNA repair (Figure 5F, Supplementary Figure 4, HC vs H). Consistently, key checkpoint transcripts were significantly reduced: p53 (G1/S checkpoint), Cdc25c (G2/M checkpoint) and Bub1 (spindle checkpoint) (Figure 6G) [28]. The downregulation of Chd4, Ezh2, Parp1, Fancl and Brca1 further supported defects in chromatin organization and DNA repair mechanisms in HC ductal cells (Figure 5G-H).

**Figure 6:**
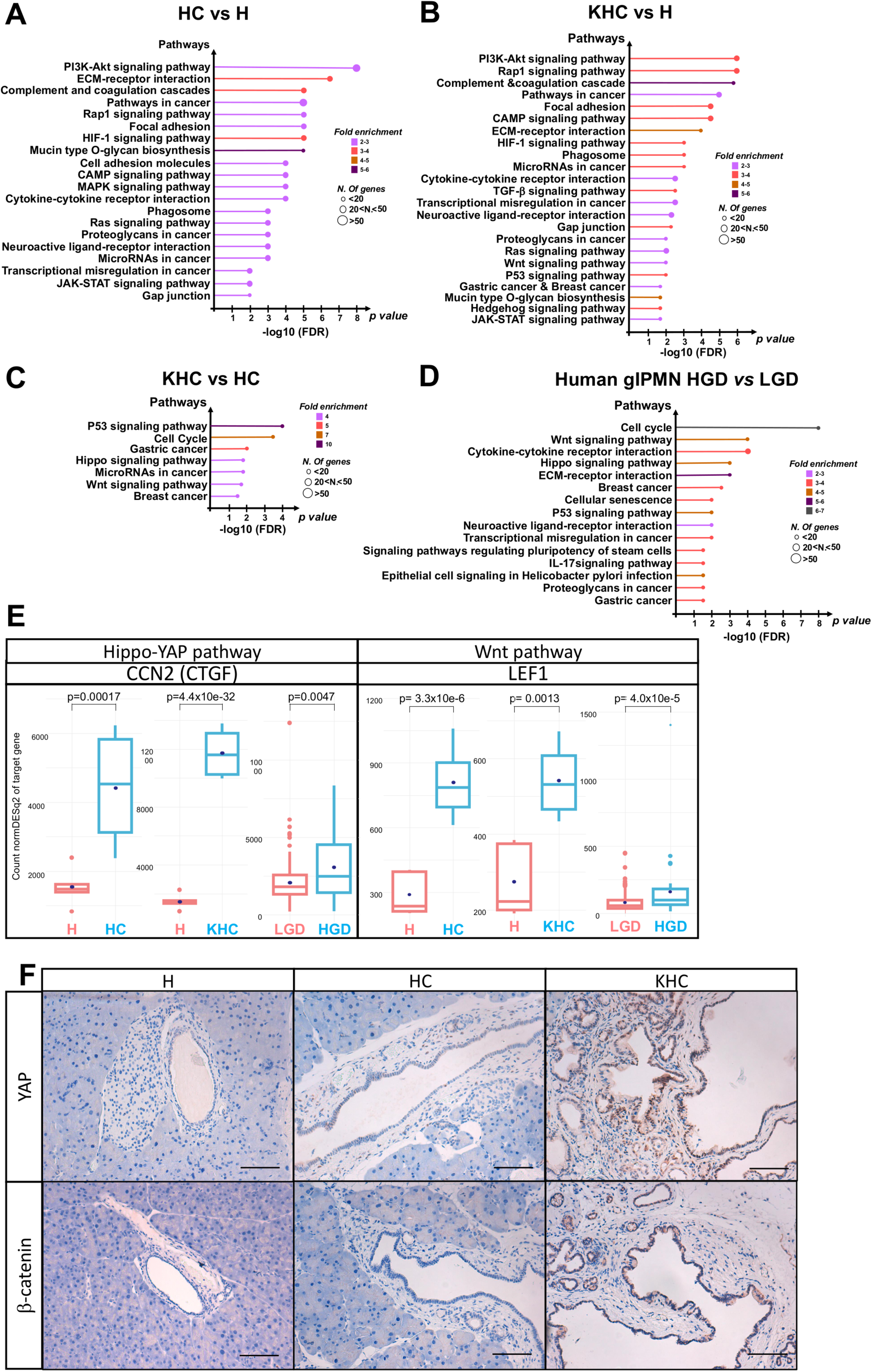
Upregulated signaling pathways in IPMN initiation and progression in mice and humans. (A) Lollipop diagram of upregulated KEGG pathways in HC compared with H. (B) Lollipop diagram of upregulated KEGG pathways in KHC compared with H. (C) Lollipop diagram of upregulated KEGG pathways in KHC compared with HC. (D) Lollipop diagram of upregulated KEGG pathways in human gastric IPMN (gIPMN), HGD compared with LGD. (E) Boxplots showing the upregulation of genes encoding effectors associated with the Hippo-YAP (CTGF/CCN2) and Wnt (LEF1) pathways in HC and KHC compared with H controls; and in human gIPMN, HGD compared with LGD. (F) Immunohistochemical staining of YAP and β-catenin showing positive labeling in KHC. Scale bar 100mm.

Importantly, KEGG revealed early activation of pro-tumor signaling in HC and KHC ducts: PI3K–AKT, RAS, RAP1, cAMP and JAK–STAT, together with ECM interactions and mucin-type O-glycan biosynthesis (Figure 6A-B; key genes in Supplementary Table 2). At the transcript level, LEF1 (Wnt) and CTGF/CCN2 (Hippo–YAP) were upregulated in HC and KHC ducts (Figure 6C). Critically, P53, Hippo-YAP and Wnt signaling were markedly upregulated in KHC versus HC cells, highlighting their central role in dysplasia development (Figure 6C; Supplementary Figure 5). Immunostaining of β-catenin and YAP1 confirmed activation of Wnt/β-catenin and YAP pathways at the protein level in KHC ducts (Figure 6F). Concordantly, human IPMN progression from LGD to HGD exhibited also upregulation of P53, Wnt and Hippo–YAP pathways (Figure 6D-E, Supplementary Figure 6).

Concomitantly, a metabolic reprogramming including Warburg effect was found in HC and KHC cells, with increased expression of Glut1/Slc2a1, Pkm2, Ldha, Acacb, Fasn and Pparγ (Supplementary Table 3) [29]. This metabolic reprogramming was also observed in human IPMN (n=179), and more pronounced in HGD (n=39) than LGD (n=140) (Figure 6D-E; Supplementary Table 3).

Collectively, these findings demonstrate that loss of *Hnf1b* disrupts primary cilium, impairs ductal function, and enables activation of pro-tumorigenic signaling, supporting an essential role in IPMN initiation.

### Increased IPMN prevalence in *HNF1B* mutation carriers

Given that *Hnf1b* deficiency predisposes to IPMN in our mouse model, we investigated whether patients with germline heterozygous *HNF1B* mutations or deletions show increased IPMN prevalence. HNF1B-related disease includes chronic renal failure, diabetes (Maturity Onset Diabetes of the Young type 5, MODY5), and frequent pancreatic hypoplasia/atrophy [30]. While renal cysts are a hallmark, pancreatic cysts, especially IPMN, remain understudied in this population.

Among 130 patients followed in the pediatric and adult nephrology departments of Necker Hospital (Paris, France), 45 (mean age 35.8±16.4 years; female n=19, 42%) had suitable kidney imaging (MRI with T2 sequences) for IPMN screening and were included. The 85 patients without imaging data were significantly younger, with a mean age of 19.8±16.9 years. Despite agenesis of the body and tail of the pancreas in 30 (67%) patients, 14 (31%) had branch-duct IPMN (Figure 7). Compared with the theoretical 5% incidence of IPMN in age-matched individuals, the significantly greater occurrence of branch-duct IPMN in patients with heterozygous *HNF1B* mutations (p=2.3×10^-8^) strongly suggests that *HNF1B* has a tumor-suppressive role in human IPMN initiation.

**Figure 7:**
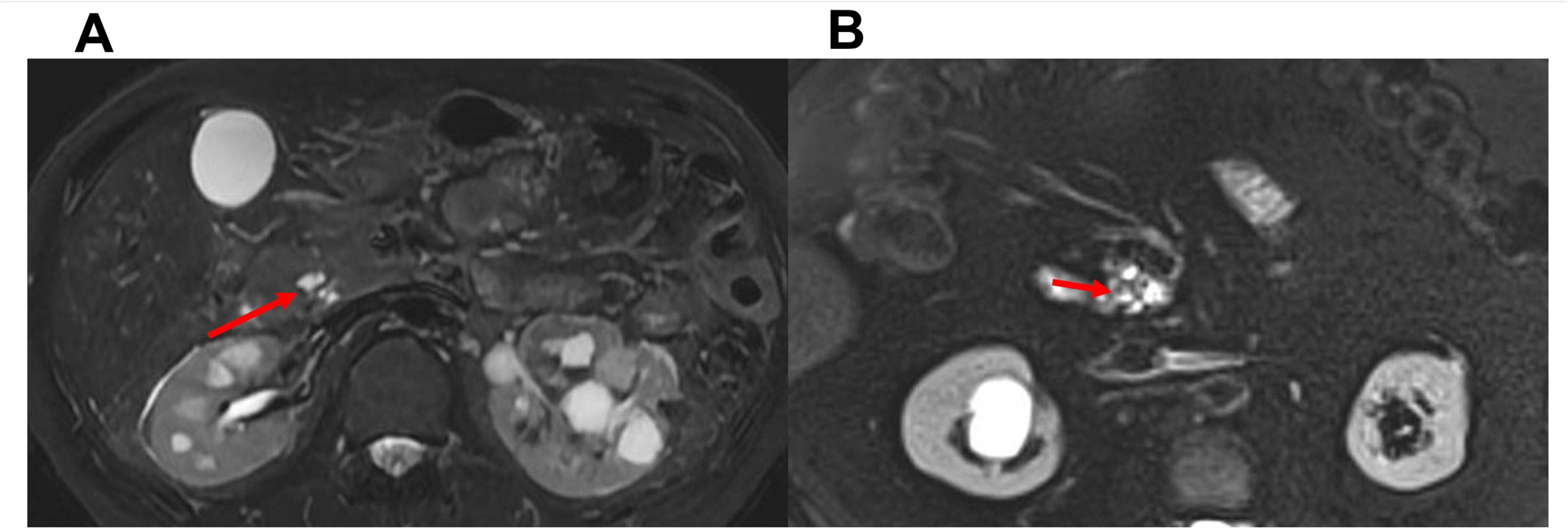
Increased IPMN prevalence in *HNF1B* mutation/deletion carriers. Axial T2-weighted MR image showing branch-duct IPMN in the pancreatic head of 2 young patients with a heterozygous germline mutation in HNF1B. The connection to the main pancreatic duct is clearly visible in (A), and mucus droplets are distinctly visible in (B).

## Discussion

In human, we found a loss of *HNF1B* expression in IPMN, as dysplasia progresses, largely via promoter methylation. In a ductal-specific mouse model, *Hnf1b* loss combined with KRAS activation causes gastric-type IPMN with primary-cilium depletion and activation of pro-tumor signaling pathways. Consistently, germline *HNF1B* carriers show a markedly increased prevalence of branch-duct IPMNs. Together, these data demonstrate a tumor-suppressive role for *HNF1B* in ductal epithelium.

Although *HNF1B* is a known susceptibility gene for pancreatic cancer, mutations have not been commonly reported in IPMNs [11,19,31,32], likely due to its exclusion from targeted sequencing panels and its complex mutational landscape, which includes over 400 known variants (ClinVar, National Center for Biotechnology Information, U.S. National Library of Medicine) [10,30,33], often within repeat-rich regions prone to replication errors [34,35]. While inactivating *HNF1B* mutations have been reported in prostate, endometrial and ovarian cancers, *HNF1B* methylation is another mechanism of gene silencing observed across various cancer types (ovarian, prostate, kidney, biliary tract, lung, colon cancers) [18,36–39]. Until now, *HNF1B* methylation had not been evaluated in pancreatic cancer or IPMN. Our data show high rates of *HNF1B* methylation in gastric and intestinal IPMN, increasing from 22% in LGD to 55% in HGD (and reaching 100% when intermediate methylation is included), supporting epigenetic silencing as a contributor to HNF1B loss in IPMN.

Mechanistically, *HNF1B* has been already identified as a tumor suppressor in various cancers, including kidney, bladder, breast, prostate, ovary, uterus, liver and colon [18,36,39–42]. Moreover, in PDAC, the loss of HNF1B correlates with poor survival and dedifferentiation [43,44]. Consistent with this, HNF1B appears to exert its tumor-suppressive function through multiple mechanisms. First, some studies have implicated dysfunction in chromatin accessibility programs, which impact epigenomic landscape [43]. In parallel, loss of the BRG1/SWI-SNF complex has also been reported as a factor triggering the onset of IPMN, according to the concept of loss of cellular identity, with the observation of “ductal regression” [22]. Besides, mitotic bookmarking by HNF1B has been described, preserving transcriptional continuity across cell division; its loss can perturb cell-cycle progression and may contribute to tumor progression [45,46]. Furthermore, other studies support a genome-surveillance role for HNF1B [41,47,48]. In the kidney, cooperation between P53, a key genome caretaker, and HNF1B has been reported, with combined loss correlating with more aggressive behaviour [42,49]. In prostate cancer, HNF1B loss has been linked to defective spindle checkpoints with reduced *MAD2L1* and *BUB1* expression (both reported as direct HNF1B targets) and consequent chromosomal mis-segregation [41,48]. In the pancreas, ChIP data from PANC1 cells identify *CDC25C* (mitotic checkpoint), *MSH2* and *BLM* (DNA repair) as HNF1B direct targets (Chip-Atlas Consortium, 2024) [42]. Consistently, in our mouse model, *Hnf1b* loss resulted in downregulation of genome surveillance pathways (mitotic checkpoint, spindle checkpoint, DNA repair) and decreased expression of several critical cell cycle regulators, including *p53*, *Bub1*, and *Ezh2*.

Loss of HNF1B appears to be an early event in IPMN tumorigenesis. Indeed, our data show that *Hnf1b*-deficient ductal cells (HC) already display a pre-tumoral changes characterized by the loss of primary cilia, the early activation of oncogenic signaling pathways (PI3K–AKT, RAS, cAMP and JAK–STAT) and metabolic reprogramming, before dysplasia onset. In human IPMNs, primary cilia loss also occurs early, already present at the stage of low-grade dysplasia [50]. Loss of the primary cilium appears to be a key initiating event, disrupting crucial signaling networks and potentially enabling additional oncogenic changes [51]. Our KHC mouse model of IPMN lesions also lacks primary epithelial cilia.

If HNF1B loss alone is not sufficient to form IPMN, it establishes a permissive ductal state from which dysplasia emerges once KRAS is activated. Indeed, KRAS mutations are major drivers of pancreatic oncogenesis in human IPMNs, present in more >95% of cases [7]; even if GNAS mutations, promoting proliferation via cAMP signaling, are more specific (60-75% of IPMNs) [7,31,52,53]. Notably, cAMP-pathway is activated in our ductal model without specific activation of GNAS. Nevertheless, IPMN onset is more complex than KRAS activation only, requiring additional genetic alterations. This highlights the need for relevant models, such as KHC. Whereas animal embryonic Cre-lox GEMMs (e.g., Pdx1-Cre, Ptf1a-Cre) activate recombination in multipotent pancreatic progenitors that later yield ductal and acinar lineages, producing lesions with mixed PanIN/IPMN features and confounding cell-of-origin, ductal-specific postnatal GEMMs (e.g., Sox9-CreER), including ours, restrict recombination to ducts and allow investigation of IPMN development. In these models, YFP lineage tracing confirms that IPMN lesions arise from ductal cells, aligning with studies identifying ducts as the IPMN cell of origin [3,20]. Across such models, dysregulated pathways reported include JAK–STAT, MAPK–EGFR, cAMP–PKA, PI3K–AKT, TGF-β, Notch and Hedgehog [3,12,13]. In our ductal model, KHC IPMN lesions show enrichment of PI3K–AKT, Rap1, C-AMP, TGF-β, Ras, Hedgehog, JAK-STAT, Wnt/β-catenin, Hippo–YAP and p53. Interestingly, the emergence of dysplasia (KHC vs HC) is accompanied by selective amplification of Wnt/β-catenin, Hippo–YAP and p53 signaling. Furthermore, we found that the same activation of the p53, Wnt/β-catenin and Hippo–YAP pathways tracks the continuum from low- to high-grade dysplasia in human surgical specimen of IPMNs. Thus, IPMN initiation is characterized by activation of several signaling pathways, but dysplastic progression is most closely linked to selective amplification of the Wnt/β-catenin, YAP and p53 pathways.

Taken together, these results establish HNF1B as a tumor-suppressive regulator of ductal epithelium. This is supported by the loss of HNF1B expression observed in human IPMN, primarily due to gene silencing through promoter methylation, and by the high prevalence of IPMNs in patients carrying constitutive HNF1B mutations/deletions. Our new GEMM model KHC supports that the loss of HNF1B is causally linked to the development of IPMNs, which is associated with cellular reprogramming, the loss of the primary cilium and the activation of pro-tumoral signaling pathways. They also highlight potential therapeutic strategies, such as restoring HNF1B expression, reestablishing ciliary function, or targeting the YAP and Wnt pathways. These insights clarify early IPMN biology and argue for monitoring individuals with HNF1B syndrome.

## Abbreviations

DAB: 3,3’-Diaminobenzidine
FDR: False Discovery Rate
FFPE: Formalin-fixed paraffin-embedded
GEMMs: genetically engineered mouse models
GOBP: Gene Ontology Biological Process
GOCC: Gene Ontology Cellular Component
GWAS: Genome-Wide Association Study
KEGG: Kyoto Encyclopedia of Genes and Genomes
HES: Hematoxylin, Eosin, and Saffron
HGD: high-grade dysplasia
IHC: immunohistochemistry
IPMN: Intraductal Papillary Mucinous Neoplasm
LGD: low-grade dysplasia
MODY: 5 maturity-onset diabetes of the young type 5
MRI: Magnetic Resonance Imaging
PanIN: Pancreatic Intraepithelial Neoplasia
PCA: Principal Component Analysis
PDAC: Pancreatic Ductal Adenocarcinoma
YFP: Yellow Fluorescent Protein.

## Acknowledgments

We acknowledge members of the animal core facility (I. Renault, S. Olivre, A. Bouhalfaia) of the Centre de Recherche sur l’Inflammation (INSERM UMR1149). This work benefited from equipment and services of the iGenSeq core facility at Institut du Cerveau et de la Moëlle épinière, Paris, for sequencing.

## Author Contributions

DL conceived and performed experiments, analyzed the data, obtained fundings and wrote the manuscript. LAM, LMD, An Ch, PN, LM, performed experiments and analyzed data. LH, BK, VR coordinated the clinical cohorts. SD performed MRI acquisition and analysis. IE shared methylation database. CP, Al Co, RN contributed to the sequencing experiment analyses. JC, AC analyzed data. CH designed the study, supervised the project, conceived and performed experiments, analyzed the data, obtained fundings and wrote the manuscript. All authors reviewed and approved the manuscript.

## Ethics statements

### Patient consent for publication

Not applicable.

### Ethics approval

This study was approved by the research Ethics Committee Institutional Review Board (IRB00006477), the Ethics Committee C2EA-121 PARIS-NORD and the French Ministry of Research (APAFiS N°34673).

