## Supplemental Material for "Loss of HNF1B drives pancreatic Intraductal Papillary and Mucinous Neoplasms (IPMN) initiation"

### ***Supplementary Material***

### ***Supplementary Material 1: Methylation analysis of HNF1B in human IPMNs***

#### **Methylation profiling methodology**

#### **Sample Collection and DNA Extraction**[9]

A cohort of 52 IPMN samples, encompassing both low-grade (n=32) and high-grade (n=20) dysplastic lesions, was obtained from resected pancreatic specimens. The histological subtypes included gastric (n=32) and intestinal (n=20) IPMN subtypes, and normal pancreatic ducts (n=19). DNA methylation profiling was performed using the Infinium Methylation EPIC BeadChip (Illumina, San Diego, USA), which covers over 850,000 CpG sites across the genome. The BeadChips were processed at the Genomics and Proteomics Core Facility of the German Cancer Research Center in Heidelberg, as described by Liffers *et al.* (2023). Sample collection and DNA extraction were carried out by the team of Irene Esposito.

#### **Infinium Methylation EPIC BeadChip Analysis**

The DNA methylation profiles were generated using the Infinium Methylation EPIC BeadChip (Illumina, San Diego, USA), which interrogates over 850,000 CpG sites across the human genome. Processing of the BeadChips was conducted at the Genomics and Proteomics Core Facility of the German Cancer Research Center in Heidelberg, as part of a collaboration.

Data processing and quality control were performed using the R Bioconductor package ChAMP (v2.14.0). IDAT files were loaded into ChAMP, and data analysis was processed with R Bioconductor package ChAMP (V.2.14.0) and Illumina Infinium HumanMethylation assay (illuminaio)[10].

#### **Probe selection and analysis of *HNF1B* methylation**

The analysis focused on 42 probes associated with the *HNF1B* gene. For the analysis, probes were specifically selected on the basis of their association with the transcriptional regulation of the *HNF1B* gene, ensuring that the probes included in the study were relevant to the gene's expression and potential epigenetic silencing mechanisms (notes for the Illumina HumanMethylationEPIC27 BeadChip /IlluminaHumanMethylationEPICanno.ilm10b4.hg19 and UCSC_RefGene_Name).

The table below summarizes the selected probes, their genomic location, their context within the *HNF1B* gene, and their probe type:

| Probe ID | Genomic Region on HNF1B | RefGene Group | Probe Type |
|---|---|---|---|
| cg04105270 | Promoter (TSS1500) | S_Shore | Type 1 |
| cg05496067 | Body | OpenSea | Type 2 |
| cg02763119 | Body | OpenSea | Type 2 |
| cg04198914 | Promoter (TSS1500) | S_Shore | Type 1 |
| cg21253112 | Body | OpenSea | Type 2 |
| cg03348978 | Body | Island | Type 2 |

**Heatmap generation**

The R code used to generate a heatmap, displaying the methylation patterns of *HNF1B* across the pancreatic ducts and IPMN can be provided upon request.

Mean beta value calculation and methylation status categorization of pancreatic ducts and IPMN[9,11,12]

Methylation levels were quantified using beta values. The following thresholds were applied to categorize the methylation status:

- Methylated**:** Beta value >0.5
- Unmethylated**:** Beta value <0.4
- Partially methylated/intermediate**:** Beta value between 0.4 and 0.5

Each IPMN sample was classified as methylated, unmethylated, or partially methylated, based on the beta values observed at the *HNF1B* locus.

**Statistical analysis**

A Fisher's exact test was utilized to compare the methylation status of *HNF1B* across different IPMN subtypes and dysplasia grades. This statistical method was chosen due to the small sample sizes in certain groups, ensuring the robustness of the analysis.

***Supplementary Material 2: RNAseq analysis***

The samples were processed at the ICM's high-throughput sequencing platform. (Institut du Cerveau et de la Moelle épinière, Hôpital Pitié-Salpêtrière Hospital, Paris, France). Gene expression quantification was carried out, resulting in the generation of gene count files for the different samples. The quality of the samples was verified after processing by checking the number of genes in the sample (greater than 20000), ensuring the absence of degradation.

RNA-seq analysis of pancreatic ductal organoids from H, HC, and KHC mice

**Process method for RNA Sequencing output:**

Reads were aligned to the reference genome (Genome Reference Consortium ensemblv107_GRCm38.75/*Mus musculus*) and assembled into transcripts using STAR[1]. Gene expression counts were obtained using FeatureCounts[2].

Three analysis platforms were utilized: IDEP 1.1[3], ShinyGO[4], and DECAFE© (*developed by Rémy Nicolle's team, Beaufils and Pignolet., 2024 ;DECAFE is licensed under CC BY-NC 4.0. To view a copy of this license, visit http://creativecommons.org/licenses/by-nc/4.0/*) to statistically analyze transcriptomic differences between the H, HC, and KHC conditions using the following methodology:

- **Principal Component Analysis (PCA):** The gene expression matrix was submitted to PCA to visualize sample clustering and identify potential outliers.
- **Venn diagram creation:** Venn diagrams were generated to graphically represent the relationships between different transcript sets, highlighting elements common to or specific to each group.
- **Heatmap generation:** Heatmaps was produced to display the gene expression patterns of the differentially expressed clusters of interest between the samples. The heatmaps was color-coded (red for high expression, blue for low expression) with dendrograms illustrating the similarity relationships between genes and samples. K-means clustering analysis was performed to group genes with similar expression patterns across the H, HC, and KHC conditions. This method facilitates the identification of distinct clusters of co-expressed genes, allowing for the detection of potential gene networks and pathways that are differentially regulated between the sample groups[5].

- **Differential Gene Expression Analysis (DESeq2):** DESeq2 was employed to analyze variations (overexpression or underexpression) between the H, HC, and KHC conditions in pairwise comparisons. DESeq2 estimates gene dispersions, fits a negative binomial model, and performs hypothesis testing using the Wald test or likelihood ratio test, with p-values adjusted using the Benjamini & Hochberg method. Results were visualized globally on a volcano plot and examined gene by gene for key genes of interest using specific gene boxplots for comparisons between HC vs. H, KHC vs. H and KHC vs HC[6].
- **Functional Analysis/ « GO-pathways » with ShinyGO:** Functional analysis was conducted using GSEA tools with the KEGG (Kyoto Encyclopedia of Genes and Genomes) database and the GO (GOBP [Gene Ontology Biological Process] and GOCC [Gene Ontology Cellular Component]) databases to interpret differentially expressed genes. This analysis identified the biological pathways and cellular processes involved, and the pathways were organized into networks using ShinyGO[7,8].

<u>RNA-seq analysis of human gastric IPMNs with Low-Grade and High-Grade Dysplasia</u>

The same RNA-seq analysis pipeline was applied to compare transcriptomic profiles between low-grade dysplasia (LGD) and high-grade dysplasia (HGD) in microdissected human gastric IPMNs obtained from surgical specimens.

- **Reads Alignment and Gene Count:** Reads were aligned to the human reference genome (ensemble 107_GRCh38.75 (*Homo sapiens* samples)) using STAR, and gene expression counts were generated with FeatureCounts.
- **Principal Component Analysis (PCA):** PCA was conducted to visualize sample clustering and detect outliers.
- **Heatmap:** Heatmaps was created to illustrate differential gene expression patterns, with K-means clustering applied to identify co-expressed gene networks that may play roles in the progression from LGD to HGD.
- **Differential Gene Expression (DESeq2):** DESeq2 was used for pairwise comparisons between LGD and HGD, identifying key differentially expressed genes. These results were visualized on volcano plots and gene-specific boxplots.
- **Functional Pathway Analysis (ShinyGO):** Functional analysis using ShinyGO, incorporating KEGG and GO databases, identified biological pathways and cellular

processes differentially regulated between LGD and HGD, shedding light on the continuum of tumorigenesis in humans.

***Supplementary Material 3: Detailed protocol of mouse pancreatic duct organoids***

**Preparation of organoid media**

- **HBSS Washing Solution:** HBSS (ThermoFisher, #14025), Complete Protease Inhibitor Cocktail (Roche, #11697498001, 1 tablet for 50 mL), and Y-27632 Rho-kinase inhibitor (Bio-techne, #1254/1, 10.5 µM).
- **Dissociation Medium:** Collagenase XI (Sigma-Aldrich, #C7657, 0.1 mg/mL), Y-27632 Rho-kinase inhibitor (Bio-techne, #1254/1, 10.5 µM).
- **HBSS Digestion Stop Solution:** HBSS (ThermoFisher, #14025), Fetal Calf Serum (ThermoFisher, #10270, 5%), Complete Protease Inhibitor Cocktail (Roche, #11697498001, 1 tablet for 50 mL), Y-27632 Rho-kinase inhibitor (Bio-techne, #1254/1, 10.5 µM).
- **Complete Culture Medium:** Advanced DMEM/F12 (ThermoFisher, #12634), HEPES (ThermoFisher, #15630, 10 mM), Penicillin/Streptomycin (ThermoFisher, #15140, 10 mM), GlutaMAX (ThermoFisher, #35050, 10 mM), Recombinant Mouse EGF (ThermoFisher, #PMG8041, 50 ng/mL), Recombinant Human FGF-10 (Sigma-Aldrich, #SRP3262, 100 ng/mL), Gastrin I (Sigma-Aldrich, #SCP0152, 0.01 µM), mNoggin (Sigma-Aldrich, #H6416, 100 ng/mL), N-acetylcysteine (Sigma-Aldrich, #A9165, 1.25 mM), Nicotinamide (Sigma-Aldrich, #72340, 10 mM), Recombinant R-Spondin (Bio-techne, #4645-RS-025, 1 µg/mL), B27 Supplement 50X Serum-Free (ThermoFisher, #17504044, 1X), Y-27632 Rho-kinase inhibitor (Bio-techne, #1254/1, 10.5 µM), Fungizone (ThermoFisher, #15290, 250 µg/mL).
- **Extracellular Matrix:** Cultrex Basement Membrane Extract (BME), Type 2 (Bio-techne, #3533-010-02).

**Pancreatic ductal organoids from mice**

Pancreatic tissues from 12-week-old H, HC, and KHC mice were collected in a 35 mm Petri dishes. Three milliliters of cold HBSS washing solution were added to the dish, and this washing step was repeated twice. The pancreatic tissues were then minced into 1.0-2.0 mm fragments using a sterile surgical scalpel. The tissue fragments were transferred to a 50 mL conical tube using a 25 mL pipette and centrifuged at 400 g for 3 minutes at 4°C. The pellet was resuspended in 3 mL of digestion medium and placed on an orbital shaker at 37°C. The digestion process was monitored every 10 minutes, with mechanical dissociation performed using a 25 mL pipette. After 40 minutes, the digestion progress was checked under a microscope. Following digestion, the samples were centrifuged for 5 minutes at 700 g at 4°C, and the pellet was resuspended in 3 mL of HBSS washing solution. This step was repeated twice before resuspending the pellet in 5 mL of HBSS to stop the digestion process. Pancreatic ducts were manually sorted under a binocular magnifier or fluorescence microscope for YFP-positive ducts. To increase purity, 2 or 3 successive sorting steps were performed. The sorted pancreatic ducts were centrifuged for 5 minutes at 700 g, and the cell pellets were resuspended in 10 µL of complete culture medium. The culture plates were pre-warmed at 37°C for several hours before use to prevent flattening of the BME domes where the organoids were embedded. The cell suspension was mixed with an appropriate volume of BME2 matrix (40 µL of matrix for 10 µL of cell suspension), and 50 µL domes were formed on a 24-well plate, followed by incubation for 30 minutes at 37°C. The domes were then covered with 450 µL of complete culture medium. The plates were incubated at 37°C in a 5% $CO_2$ humidified incubator. The organoid cultures were observed daily using an inverted LEICA DMIRB fluorescence microscope. The medium was changed every 2-3 days and 500 µL of pre-warmed (37°C) complete culture medium was added dropwise to the well. The organoids were grown for 7 days to allow the selection and amplification of ductal organoids. On day 7, a portion of the organoids was fixed and embedded in paraffin for histological analysis (Sox9 immunohistochemistry), while the remaining part was used for RNA extraction.

**Inclusion for Immunohistochemistry**

The culture medium was replaced with 500 µL of 4% formaldehyde (buffered, pH=7) containing erythrosine (0.3%), a pink dye used to mark and visualize the organoids. The organoids were fixed for 2 hours at room temperature, then centrifuged for 5 minutes at 800 g and washed with 500 µL of PBS (ThermoFisher, #14190). The pellet was resuspended in 150 µL of melted Type VII agarose (Sigma, #A0701, 0.4 g/10 mL) and placed on 200 µL of solidified Type VII agarose. The agarose was allowed to polymerize on ice, forming a cylinder containing the organoids, which was then sent to pathology for paraffin embedding following the same procedure used for mouse pancreatic tissues. Sox9 immunostaining was performed on the organoids to verify their ductal nature.

### RNA Extraction

RNA extraction was performed using the RNeasy Micro Kit (Qiagen, #74004), allowing for the lysis of ductal organoids, RNA purification, washing, and elution. No carrier RNA was used to avoid distorting the RNA-seq analysis results. The extracted RNA was quantified and stored at -80°C prior to RNA-seq analysis.

### Supplementary Figures and Tables

#### Supplementary Figure 1: Mean CpG methylation of HNF1B probes in human intestinal or gastric IPMN

(A) Methylation of Intestinal IPMN: Mean CpG methylation of *HNF1B* probes known to modulate *HNF1B* expression in human ducts (n=19), LG-iIPMN (n=8), HG-iIPMN (n=12). (unmethylated: mean β-value <0.4; intermediate: mean β-value >0.4 and <0.5; methylated: mean β-value >0.5).
(B) Methylation of gastric IPMN: Mean CpG methylation of *HNF1B* probes known to modulate *HNF1B* expression in human ducts (n=19), LG-gIPMN (n=24), HG-gIPMN (n=8). (unmethylated: mean β-value <0.4; intermediate: mean β-value >0.4 and <0.5; methylated: mean β-value >0.5).
HG-IPMN (high grade-IPMN), LG-IPMN (low grade-IPMN), IPMN (intraductal papillary mucinous neoplasms), iIPMN (intestinal IPMN), gIPMN (gastric IPMN).

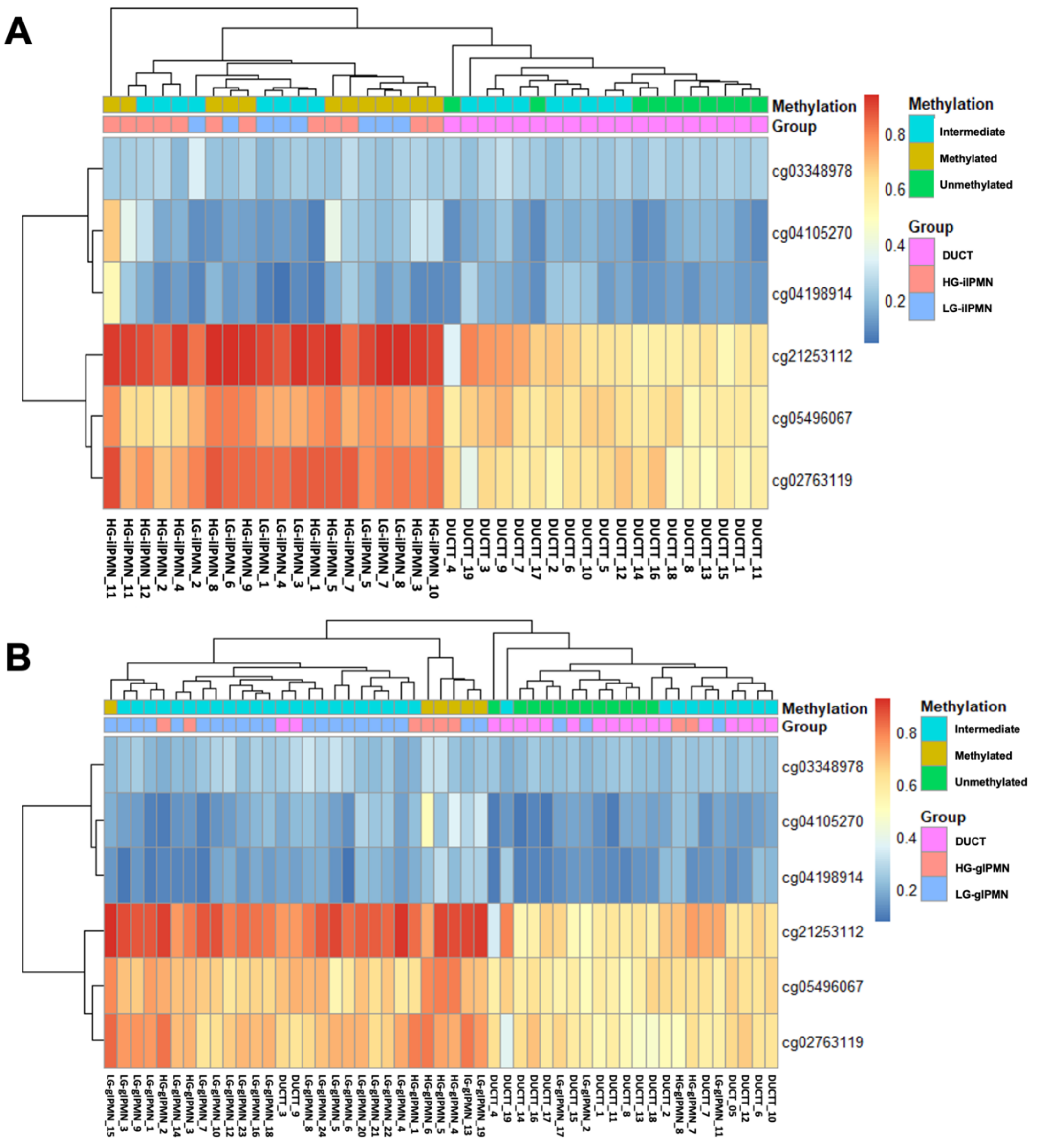

**Supplementary Figure 2: Gastric subtype of murine KHC IPMN**

(A) HES staining of human gastric IPMN and KHC mice IPMN showing histological similarities, with columnar-shaped cells and mucinous neoplasms. (B) MUC1 and MUC2 IHC of KHC mice IPMN, with mouse stomach as a positive control for MUC1 immunostaining and mouse intestine as a positive control for MUC2 immunostaining. Scale bar, 100 µm.

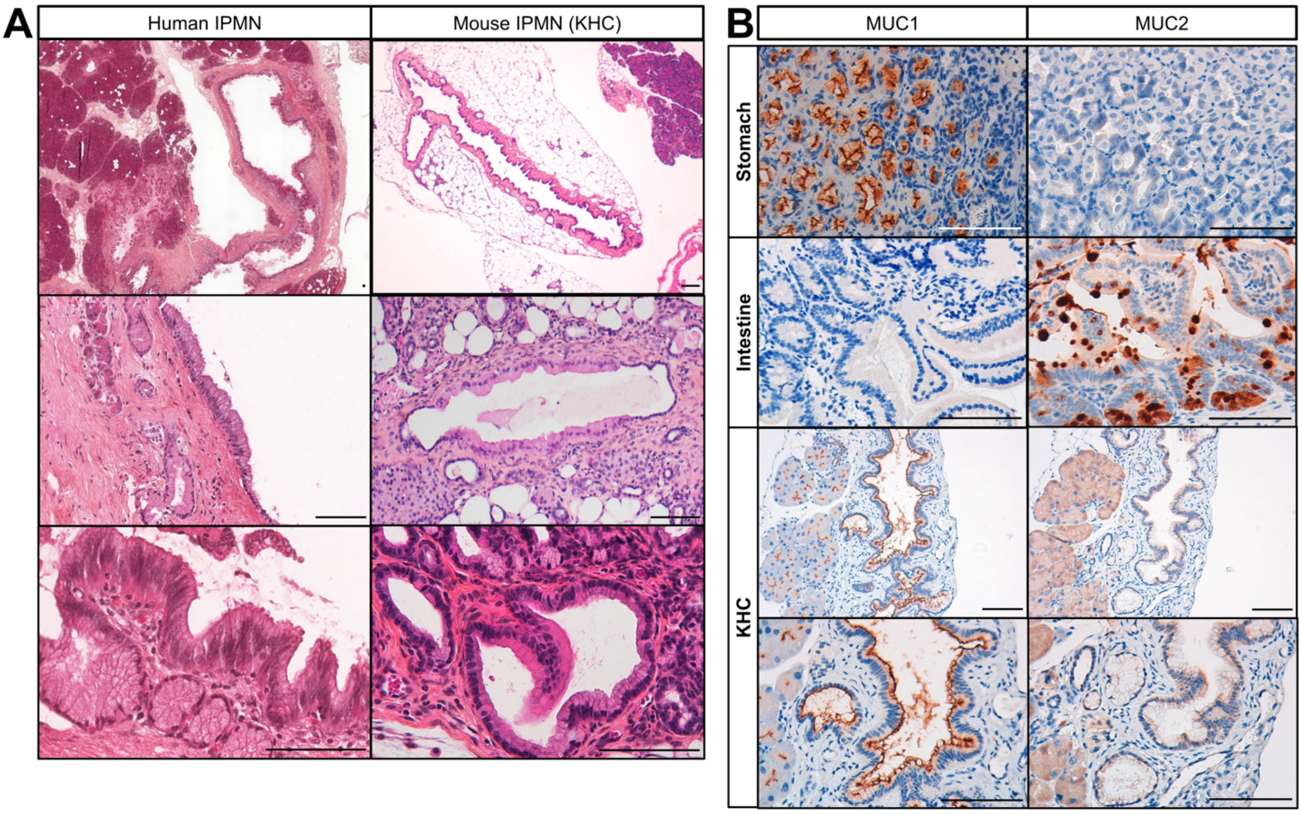

**Supplementary Figure 3: Morphology of H, HC and KHC pancreatic ductal organoids.**

(A) Increases in the number and size of H, HC, and KHC pancreatic ductal organoids. Scale bar = 100 μm. (B) Bright field and green fluorescence images of organoid wells from H, HC, and KHC mice on day 7 (D7) showing "typical" round organoids. (C) Bright field and green fluorescence images of organoid wells from HC, and KHC mice on D7 showing "tubular" organoids. (D) SOX9 immunostaining confirmed the ductal identity of H, HC and KHC organoids. Genotypes of the mice: "H" Hnf1b$^{fl/fl}$;R26R$^{YFP}$, "HC" Sox9-creER;Hnf1b$^{fl/fl}$;R26R$^{YFP}$, "KHC" Sox9-creER;LSL-Kras$^{G12D}$;Hnf1b$^{fl/fl}$; R26R$^{YFP}$.
Scale bar = 50 μm.

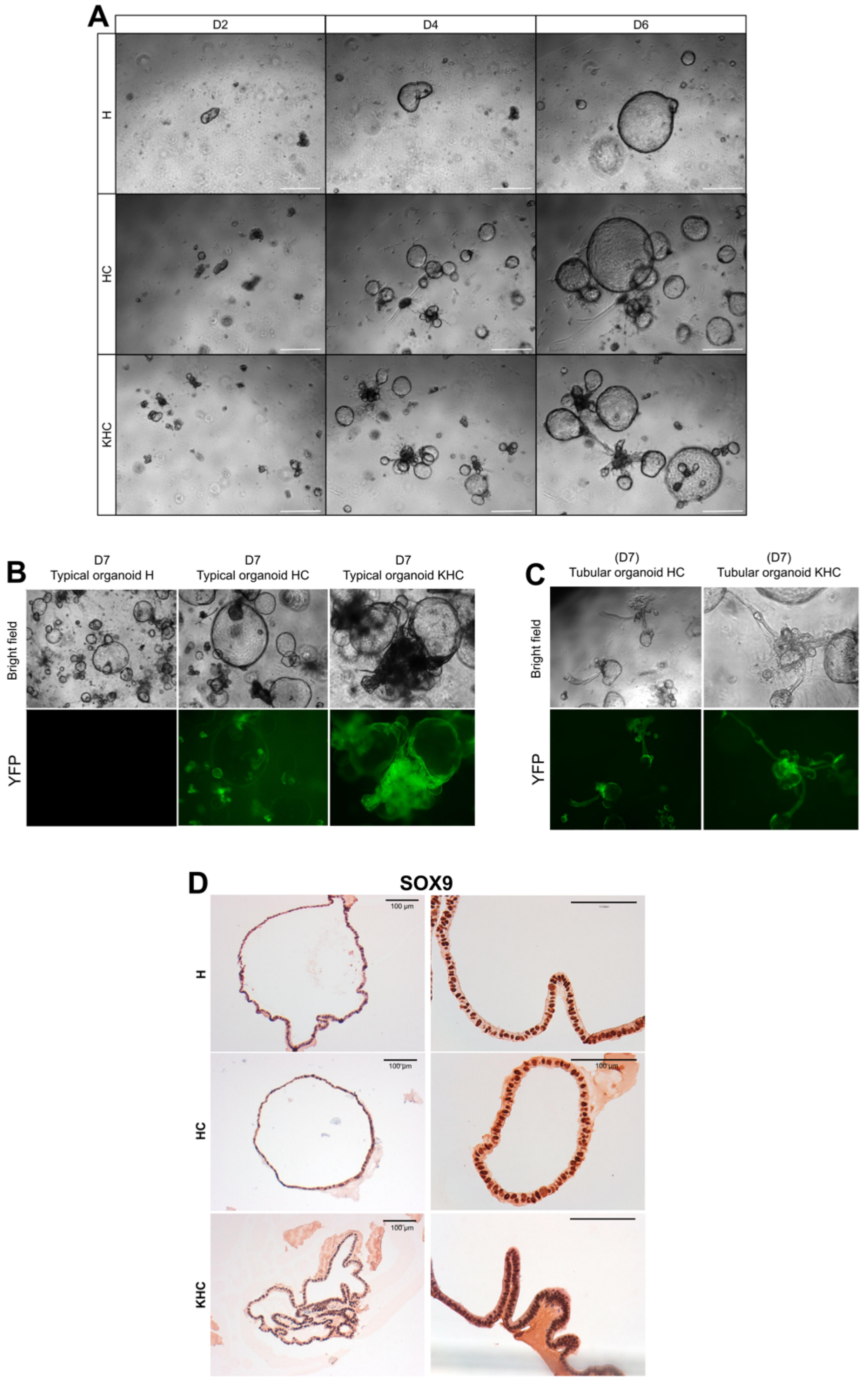

**Supplementary Figure 4: Network diagram highlighting key downregulated biological processes in HC ductal cells**

This network diagram illustrates the major biological processes that are significantly downregulated in *Hnf1b*-deficient pancreatic ductal cells compared with control ductal cells. The nodes represent individual Gene Ontology (GO) terms, with the size of each node corresponding to the significance of the downregulation. The nodes are grouped into broader functional categories, including: DNA repair pathways (red): Processes related to the recognition and repair of DNA damage; Mitotic spindle (brown): Pathways involved in spindle assembly and chromosome segregation during cell division; Cilium structure and function (blue): Pathways associated with the formation and maintenance of cilia; Regulation of chromatin structure and function (orange): Processes that modulate chromatin organization and gene expression; Cell division and its checkpoints (green): Pathways critical for cell cycle regulation and checkpoint control. The connections between nodes reflect the functional interdependencies and coordination between these biological processes, highlighting their collective downregulation in the context of HC ductal cells compared with control ductal cells.

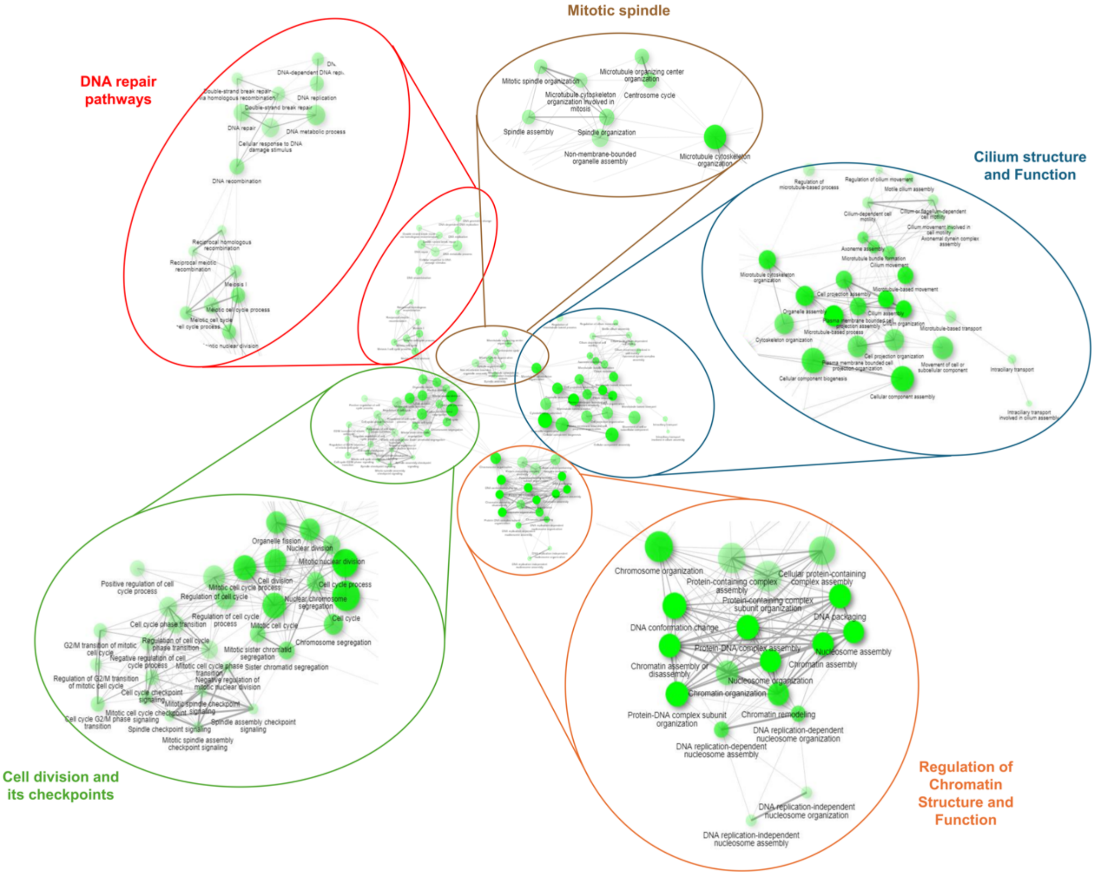

**Supplementary Figure 5: Boxplots of Hippo-YAP and Wnt pathway genes showing the upregulation of these pathways in KHC compared with HC.**

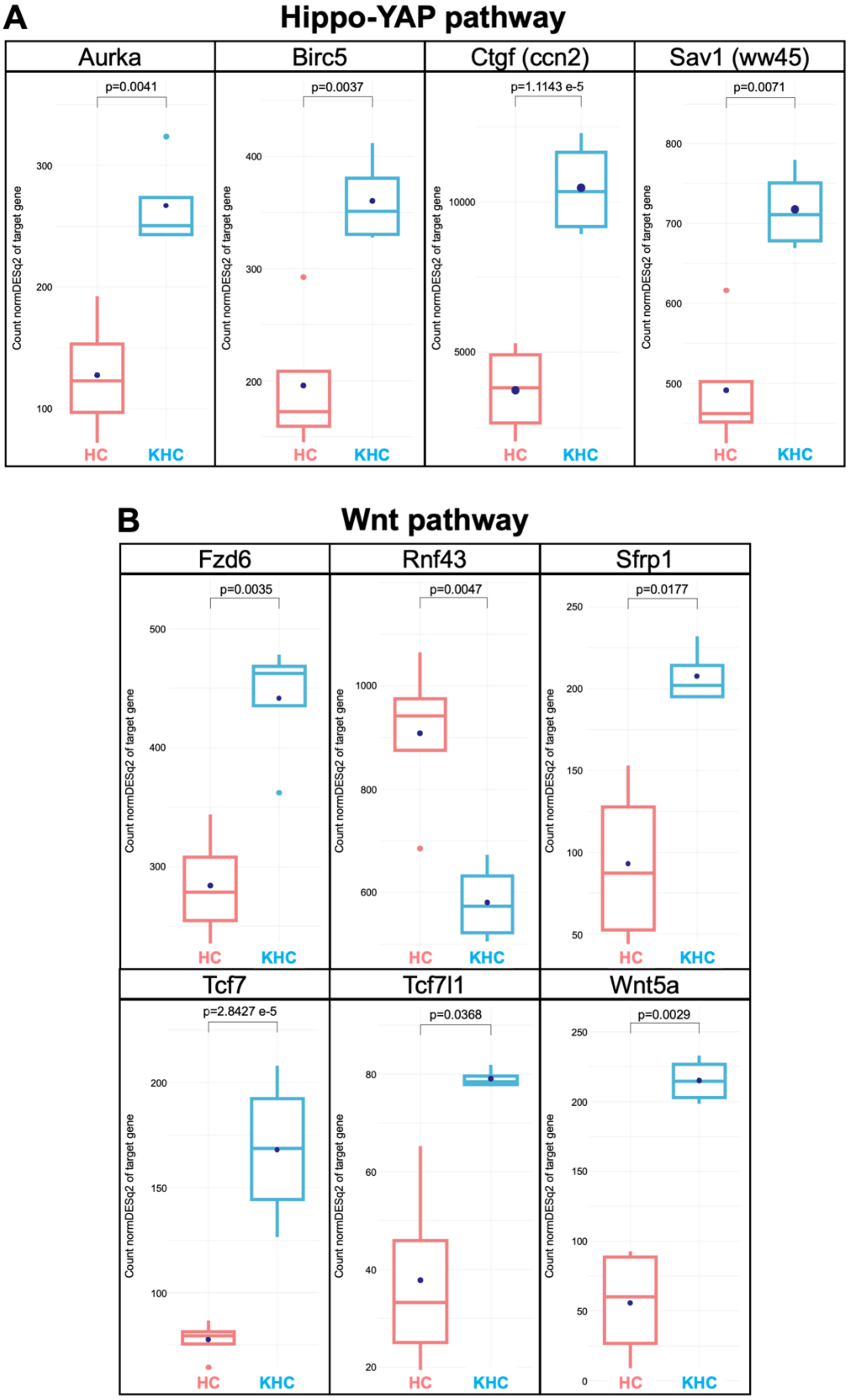

**Supplementary Figure 6: RNAseq profile of human gastric IPMN with High-Grade Dysplasia (HGD) and Low-Grade Dysplasia (LGD)**

(A) Principal component analysis (PCA) of gene expression data (LGD – red dots: n=140 / HGD – blue triangles: n=39). (B) Diagram showing the number of differentially expressed genes between HGD and LGD IPMN: 2618 downregulated / 722 upregulated genes. (C) Heatmap illustrating the clustering of IPMN samples into LGD and HGD, with notable heterogeneity within these two groups (500 genes with the most variation) (red samples: LGD / blue samples: HGD). (D) Volcano plot of differentially expressed genes between HGD and LGD in human gastric IPMN. Note that PPY gene is marked with a star as it has been moved for clarity (actual position: log2FoldChange: -21 / -log10(padj): 86). (E) Boxplot of key upregulated genes in HGD lesions, previously reported in pancreatic carcinogenesis.

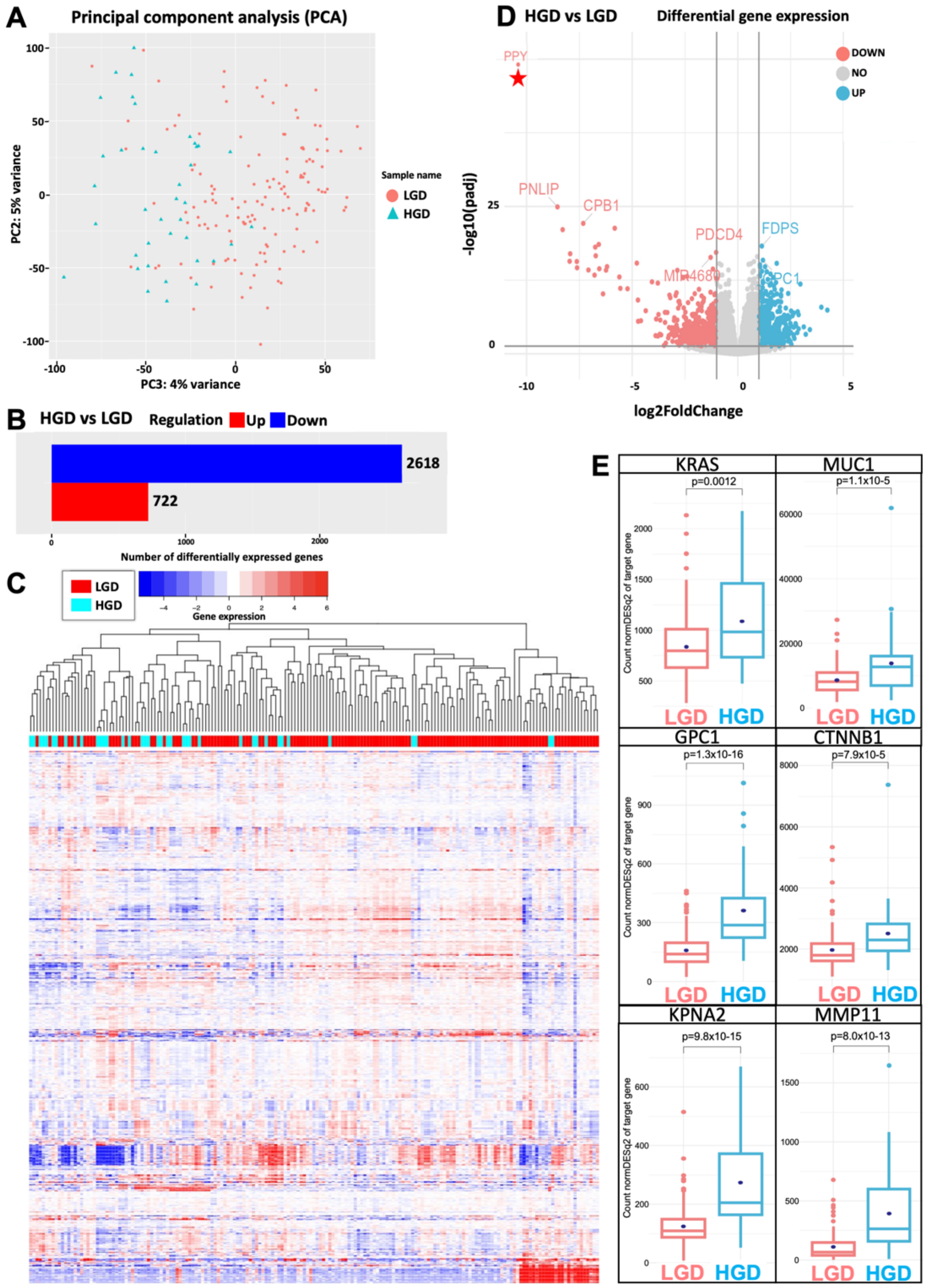

**Supplementary Table 1: List of antibodies**

| **Antibody** | **Reference** | **Host** | **Dilution** |
|---|---|---|---|
| Anti-GFP | antibody Chicken IgY, GFP-1010 ; Aveslabs | chicken | 1/200 |
| Claudin 18 | 700178 ; Thermofisher Scientific | rabbit | 1/250 |
| HNF1B | HPA002083 ; MERCK | rabbit | 1/700 |
| Hoechst | 94403; MERCK | | 1/1000 |
| Ki67 | AB15580 ; Abcam | rabbit | 1/300 |
| MUC1 | AB15481 ; Abcam | rabbit | 1/200 |
| MUC2 | AB97386 ; Abcam | rabbit | 1/200 |
| Secondary antibody IF | AlexaFluor$^{TM}$ 488 Goat anti-chicken IgY Invitrogen A11039 | goat | 1/500 |
| SOX9 | AB5535 ; Sigma Aldrich | rabbit | 1/600 |

### Supplementary Table 2: Differential gene expression in HC and KHC ducts compared with H ducts involved in molecular signaling pathways and the primary cilium

| Signaling pathways | Gene | HC vs H (mouse) | | KHC vs H (mouse) | | HGD vs LGD (human) | | Function |
|---|---|---|---|---|---|---|---|---|
| | | significance | p | significance | p | significance | p | |
| Hnf1B | Hnf1b | down | 3.5e-3 | down | 2.3e-7 | x | 0.11 | |
| known target gene of Hnf1B | Hnf6 (onecut1) | x | 6.4e-1 | down | 9.68e-19 | x | 0.67 | |
| | FGFR4 | down | 3.3e-2 | down | 1.98e-7 | x | 0.67 | |
| | spp1 | down | 2.7e-3 | down | 2.71e-7 | x | 0.29 | |
| | Cys1 | down | 3.94e-6 | down | 4.5e-11 | x | 0.49 | |
| | kif12 | down | 8.57e-16 | down | 1.13e-38 | down | 1.1e-2 | |
| | pkhd1 | down | 7.3e-4 | down | 2.63e-15 | x | 0.99 | |
| function and structure of primary cilia in epithelial pancreatic cells | ift74 | down | 2.2e-8 | down | 2.0e-3 | down | 4.9e-3 | Formation of primary cilia |
| | CPAP/cenpj | down | 3.2e-4 | down | 1.4e-2 | up | 2.7e-2 | Formation of primary cilia |
| | kif3a | down | 3.1e-2 | down | 5.3e-3 | x | 0.12 | Function of cilia |
| | BBS 1 | down | 8.0e-3 | down | 3.2e-3 | down | 1.9e-2 | Formation and function of primary cilia |
| | ALMS1 | down | 3.7e-2 | down | 3.4e-2 | x | 0.53 | Formation and function of primary cilia |
| | TCTN2 | down | 3.8e-2 | down | 1.9e-3 | x | 0.43 | Formation of primary cilia |
| | arl13b | down | 1.3e-3 | down | 5.6e-2 | up | 0.02 | Formation and function of primary cilia |
| | rsph9 | down | 1.5e-4 | down | 1.84e-6 | x | 0.79 | Structural of primary cilia |
| | B9d1 | down | 1.85e-5 | down | 1.2e-2 | x | 0.32 | Formation and function of primary cilia |
| | oscp1 | down | 7.1e-8 | down | 3.0e-6 | x | 0.33 | Formation and function of primary cilia |
| | TTC21B | down | 1.8e-2 | x | 1.9e-1 | down | 3.0e-4 | Essential for cilia biogenesis and ciliary signaling |
| | PKD1 | down | 7.3e-4 | down | 2.63e-15 | x | 0.99 | Cellular signaling and maintenance of primary cilia |
| | ift88 | down | 3.5e-2 | x | 1.1e-1 | x | 0,62 | Assembly and maintenance of cilia through intraflagellar transport |
| | CEP290 | down | 4.8e-2 | x | 7.5e-1 | down | 8.3e-4 | Assembly and stability of primary cilia |
| | CC2D2A | down | 7.3e-9 | down | 7.6e-5 | x | 0.2 | Required for the development and maintenance of primary cilia |
| | OFD1 | down | 2.9e-3 | down | 1.1e-2 | down | 3.8e-4 | Involved in cilia biogenesis and ciliopathy syndromes |
| JAK-STAT pathway | pdgfra | up | 1.5e-2 | up | 3.6e-2 | x | 0.77 | Signaling Pathway activating Regulator Genes |
| | IL2rg | up | 1.5e-2 | up | <0.05 | x | 0.68 | Signaling Pathway activating Regulator Genes |
| | Prlr | up | 1.2e-3 | up | 3.8e-4 | x | 0.52 | Signaling Pathway activating Regulator Genes |
| | JAK3 | up | 2.0e-2 | up | 3.6e-4 | x | 0.15 | Signaling Pathway activating Regulator Genes |
| | Aox1 | up | 1.4e-3 | up | 6.96e-53 | x | 0.61 | Effector Genes |
| | Aox3 | up | 1.32e-16 | up | 1.76e-26 | down | 1.4e-1 | Effector Genes |
| | CCND2 | up | 5.2e-3 | up | 5.3e-7 | x | 0.14 | Effector Genes |
| | CCND3 | up | 3.1e-3 | up | 2.1e-3 | x | 0.11 | Effector Genes |
| MAPK pathway | Raf a | x | NS | x | NS | x | 0.47 | Signaling Pathway activating Regulator Genes |
| | Kras | x | 9.0e-2 | x | 4.4e-1 | up | 1.2e-3 | Signaling Pathway activating Regulator Genes |
| | MAP2K1 | up | 8.6e-3 | x | NS | up | 1.03e-7 | Signaling Pathway activating Regulator Genes |
| | MAPK3 | up | 2.0e-2 | x | NS | down | 3.4e-2 | Signaling Pathway activating Regulator Genes |
| | cacng7 | up | 1.42e-5 | up | 7.8e-3 | down | 0.01 | Signaling Pathway activating Regulator Genes |
| | ntrk3 | up | 2.6e-3 | up | 3.0e-3 | x | 0.39 | Signaling Pathway activating Regulator Genes |
| | Rasgrf1 | up | 1.1e-2 | up | 8.0e-4 | x | 0.85 | Signaling Pathway activating Regulator Genes |
| RAS pathay | Rasgrf1 | up | 4.0e-3 | up | 7.9e-3 | x | 0.85 | Signaling Modulator Genes |
| | Rasgrf2 | up | 1.1e-3 | up | <0.05 | x | 0.56 | Signaling Modulator Genes |
| | Rac2 | up | 5.3e-3 | up | <0.05 | x | 0.15 | Signaling Modulator Genes |
| | Rassf 5 | up | 3.6e-2 | up | 2.6e-3 | down | 3.1e-2 | Effector Genes |
| | Grin 1 | up | 5.48e-6 | up | 7.25e-5 | x | 0.35 | Signaling Pathway activating Regulator Genes |
| TGF-Beta pathway | TGF-BRII | up | 4.7e-4 | up | 1.4e-3 | x | 9.0e-2 | Signaling Pathway activating Regulator Genes |
| | BMP2 | up | 1.04e-8 | x | NS | x | 0.57 | Signaling Pathway activating Regulator Genes |
| | Inhbb | x | 5.6e-2 | up | 2.3e-2 | x | 0.25 | Signaling Pathway activating Regulator Genes |
| | ACVr1b | down | 2.6e-2 | up | 3.7e-4 | down | 8,2xe-9 | Signaling Pathway activating Regulator Genes |
| | Tfr 2 | up | <0.05 | up | <0.05 | x | 0.11 | Signaling Pathway activating Regulator Genes |
| | Cdkn2b (p15) | up | 2.6e-2 | up | 4.13e-6 | x | 0.79 | Effector Genes |
| | Id | up | 2.1e-3 | up | 6.2e-3 | x | 0.43 | Effector Genes |
| | Rgma | up | 5.7e-2 | up | 6.55e-7 | x | 0.94 | Signaling Pathway activating Regulator Genes |
| | smurf1 | up | 2.0e-2 | up | 5.8e-7 | x | 0.19 | Signaling Pathway inhibiting Regulator Genes |
| Hippo-YAP pathway | CTGF | up | 1.7e-4 | up | 4.42e-32 | up | 4.8e-3 | Effector Genes |
| | Lamc2 | up | 2.84e-5 | up | 6.1e-3 | up | 2.4e-4 | Effector Genes |
| | ajuba | down | 9.91e-5 | down | 1.1e-2 | up | 3.1e-3 | Signaling Pathway inhibiting Regulator Genes |
| | Itg b2 | up | 4.2e-3 | up | 4.17e-5 | NS | 0.23 | Effector Genes |
| cAMP-PKA | Pde 10a | up | 1.3e-3 | up | 4.0e-2 | up | 9.6e-4 | |
| | Rac 1 | x | 2.9e-1 | up | 5.8e-2 | x | 0.99 | |
| | Rac 2 | up | 5.0e-3 | up | <0.05 | x | 0.16 | |
| Wnt/B-catenin pathway | frizzled 7 | up | 9.0e-3 | up | 3.3e-7 | x | 0.15 | Signaling Pathway activating Regulator Genes |
| | frizzled 1 | x | 9.0e-2 | up | 6.2e-3 | x | 0.96 | Signaling Pathway activating Regulator Genes |
| | frizzled 10 | x | 9.0e-1 | x | 9.0e-1 | up | 2.2e-5 | Signaling Pathway activating Regulator Genes |
| | tcf7 | x | 1.5e-1 | x | 1.7e-1 | up | 8.0e-4 | Signal Transduction Pathway Genes |
| | Wisp (ccn4) | x | 4.5e-1 | x | 9.9e-1 | up | 3.2e-4 | Effector Genes |
| | Fra-1 (FosL1) | x | 9.9e-2 | x | 7.6e-1 | up | 4.2e-3 | Effector Genes |
| | lrp 5 | up | 2.7e-2 | up | 6.8e-2 | x | 0.75 | Signaling Pathway activating Regulator Genes |
| | TCF 4 | up | 3.25e-6 | up | 1.3e-3 | x | 0.39 | Signal Transduction Pathway Genes |
| | axin 2 | up | 1.6e-3 | up | 1.8e-5 | x | 0.83 | Effector Genes |
| | Porcn | up | 1.2e-9 | up | 3.9e-2 | x | 0.36 | Signaling Pathway activating Regulator Genes |
| Wnt planar cell polarity | Ror 2 | up | 6.0e-2 | up | <0.05 | x | 0.95 | Signaling Pathway activating Regulator Genes |
| | Prickle 2 | up | 3.2e-2 | x | 3.6e-1 | x | 0.15 | Signaling Modulator Genes |
| | Rac 2 | up | 5.0e-3 | up | <0.05 | x | 0.16 | Signaling Modulator Genes |
| Wnt Ca2+ pathway | Wnt 5b | up | 7.0e-2 | up | 2.03e-10 | x | 0.96 | Signaling Pathway activating Regulator Genes |
| | Plce 1 | up | 3.3e-3 | x | 1.1e-1 | x | 0.15 | Signaling Modulator Genes |
| | Camk2d | up | 2.0e-3 | up | 8.9e-5 | down | 1.3e-6 | Signaling Modulator Genes |
| | Camk2b | up | 4.7e-6 | up | 2.6e-3 | x | 0.13 | Signaling Modulator Genes |
| PI3-AKT /mTOR | Rictor (mTorc2) | up | 6.5e-5 | up | 1.5e-1 | x | 0.32 | Signaling Modulator Genes |
| | Sgk1 | up | 3.9e-3 | up | 1.1e-4 | x | 0.7 | Effector Genes |
| | Itga 11 | up | 5.52e-9 | up | 1.16e-12 | up | 3.2e-3 | Signaling Pathway activating Regulator Genes |
| | Itga 7 | up | 1.4e-2 | up | 1.0e-3 | up | 5.7e-2 | Signaling Pathway activating Regulator Genes |
| | Irs1 | up | 1.25e-5 | x | 6.5e-1 | x | 0.15 | Signaling Modulator Genes |
| | pi3Kr1 | up | 3.28e-5 | up | 5.15e-18 | x | 0.34 | Signaling Pathway activating Regulator Genes |
| RAP1 pathway | Itgam | up | 1.3e-4 | up | 2.2e-4 | x | 9.5e-2 | Effector Genes |
| | Itgb2 | up | 4.2e-3 | up | 4.17e-5 | x | 0.23 | Effector Genes |
| | APBB1ip (RIAM) | up | 4.9e-4 | up | 2.9e-2 | up | 3.4e-2 | Signaling Modulator Genes |
| | Fyb (Adap) | up | 5.2e-5 | up | 7.7e-6 | x | 0.39 | Signaling Modulator Genes |
| | Lcp 2 (slp76) | up | 5.1e-4 | up | 1.7e-2 | x | 0.48 | Signaling Modulator Genes |
| Hh pathway | Beta arrestin | up | 1.0e-4 | up | 4.3e-3 | x | 0.37 | Signaling Modulator Genes |
| | RPGRIP1L | down | 1.7e-3 | up | 1.3e-2 | x | 0.65 | Signaling Pathway activating Regulator Genes |
| | Gli3 | up | 2.5e-3 | up | 2.05e-6 | x | 0.33 | Signal Transduction Pathway Genes |
| | Gli2 | up | 3.3e-2 | up | 5.1e-2 | up | 3.8e-4 | Signal Transduction Pathway Genes |
| | Ihh | up | 1.92e-5 | x | 9.0e-2 | x | 0.89 | Signaling Pathway activating Regulator Genes |
| | shh | up | <0.05 | x | NS | x | 0.12 | Signaling Pathway activating Regulator Genes |
| NOTCH pathway | Hey 2 | up | <0.05 | x | NS | x | 0.74 | Effector Genes |
| | Notch 3 | up | 9.4e-6 | up | 4.06e-6 | up | 2.2xe-10 | Signaling Pathway activating Regulator Genes |
| | Dll 4 | up | 4.0e-7 | up | 4.2e-2 | x | 0.94 | Signaling Pathway activating Regulator Genes |
| | numbl | up | 1.1e-2 | up | 4.2e-2 | up | 7.9e-3 | Signaling Pathway inhibiting Regulator Genes |

### Supplementary Table 3: Metabolic reprogramming of HC and KHC ductal cells

| Up or downregulated / p | HC vs H | KHC vs H | Human (HGD vs LGD) |
|---|---|---|---|
| **Lipid metabolism** | | | |
| Hmgcl | **Up 0.000079** | 0.89 | **Down 0.00018** |
| ALCAM | **Up 0.00088** | **Up 0.01** | 0.22 |
| EPHA2 | **Up 0.12** | **Up 9.7x10-6** | **Up 0.02** |
| FLNB | 0.16 | 0.81 | 0.57 |
| PLAU | 0.67 | 0.35 | **Up 0.0012** |
| SEMA3C | 0.68 | 0.91 | 0.52 |
| SQLE | 0.61 | 0.69 | **Up 4.6x10-7** |
| Lipe hormone-sensitive lipase (hsl) | **Up 4.4 x10-7** | **Up 1.23x10-10** | 0.77 |
| ACACB (acetylCoa carboxylase B) | **Up 1.62x10-7** | **Up 6.33x10-10** | **Down 2.35x10-5** |
| Plin1 | 0.89 | 0.71 | 0.10 |
| PNPLA2 | 0.94 | 0.94 | 0.29 |
| Fasn (Fatty Acid Synthase) | **Up 0.088** | **Up 0.0011** | **Up 7.7x10-9** |
| PPARγ (Peroxisome Proliferator-Activated Receptor Gamma) | **Up 2.3x10-9** | **Up 0.000022** | 0.37 |
| ACLY | 0.80 | 0.69 | **Up 0.04** |
| SREBF1 | 0.29 | 0.17 | **Up 0.00012** |
| Scd1 (Stearoyl-CoA Desaturase 1) | **Up 4.5x10-6** | **Up 8.1x10-10** | **Up 6.3x10-12** |
| ACACA (Acetyl-CoA Carboxylase Alpha) | 0.18 | 0.99 | **Up 6.5x10-7** |
| FADS2 (Fatty Acid Desaturase 2) | 0.47 | 0.92 | **Up 2.3x10-12** |
| **Glucid metabolism** | | | |
| IGFBP 3 | **Up 0.0019** | **Up 0.000074** | **Up 0.00056** |
| IGF1 | **Up 0.015** | **Up 0.036** | 0.44 |
| IRS1 | **Up 1.26x10-5** | **Up 6.05x10-15** | 0.15 |
| Pgm1 (PYG phosphoglucomutase) | **Up 0.0034** | **Up 0.019** | **Up 0.07** |
| Gpi1 (Glucose-6-Phosphate Isomerase 1**)** | **Up 2.57x10-9** | **Up 0.0013** | **Up 0.00033** |
| EPAC2/Rapgef4 | **Up 0.018** | 0.81 | 0.56 |
| SLC2A1 / GLUT 1 | **Up 0.0071** | 0.25 | **Up 0.00039** |
| PFKFB3 | **Up 0.00022** | 0.11 | 0.80 |
| HK2 (Hexokinase 2) | **Up 0.05** | 0.36 | **Up 0.0012** |
| PKM2 (Pyruvate Kinase M) | **Up 0.0095** | **Up 0.019** | **Up 4.38x10-14** |
| PFKP/PFK1 Phosphofructokinase-1 | **Up 6.75 x10-6** | **Up 0.09** | **Up 2.6x10-5** |
| LDHA (Lactate Dehydrogenase A) | **Up 0.038** | **Up 0.032** | **Up 1.79x10-5** |
| G6pd | 0.63 | 0.92 | **Up 0.00039** |
| PGK1 | **Up 0.000023** | **Up 0.0021** | **Up 3.8x10-7** |
| ENO1 (Enolase 1) | **Up 0.17** | **Up 0.0038** | **Up 1.4x10-12** |
| ALDOA (Aldolase A) | **Up 0.0085** | **Up 0.043** | **Up 2.6x10-5** |
| TPI1 (Triosephosphate Isomerase 1) | **Up 0.027** | **Up 0.048** | **Up 1.4x10-6** |
| GAPDH (Glyceraldehyde-3-Phosphate Dehydrogenase) | **Up 0.0084** | **Up 0.016** | **Up 4.4x10-10** |
| PDK1 (Pyruvate Dehydrogenase Kinase 1) | **Up 0.02** | **Up 0.05** | 0.29 |
